# A Neuroimmune Feedforward Circuit Linking REM Sleep with Stress-Related Behavioral Susceptibility

**DOI:** 10.64898/2026.06.04.730038

**Authors:** Binghao Zhao, Shuxian Yang, Yujie Xie, Lina Wang, Qiuqiong Huang, Jingfei Li, Xiyu Zhao, Xiao Yu, Helmut Kettenmann, Yu-Ting Tseng, Liping Wang

## Abstract

Rapid eye movement (REM) sleep is strongly associated with stress susceptibility, yet the underlying neuroimmune mechanisms remain unclear, limiting therapeutic targeting. Here, we identify a pathway by which chronic stress increases splenic nerve activity, promoting recruitment of circulating monocytes to the brain through the choroid plexus. In the external globus pallidus (GPe), recruited monocytes drive regionally restricted microglial remodeling toward a pro-inflammatory state. These microglia, in turn, potentiate the sustained excitability of parvalbumin-positive (PV⁺) neurons via IL6-gp130 signaling. Functionally, activity of GPe PV⁺ neurons during REM sleep, but not wakefulness, bidirectionally regulates anxiety-like and defensive behaviors, establishing REM sleep as a critical state through which this neuroimmune circuit governs behavioral susceptibility. Together, these findings establish a neuroimmune feedforward mechanism linking peripheral immune activation to neuronal dysregulation underlying behavioral abnormalities, and position REM sleep as a predictive and actionable physiological entry point for mitigating stress-related emotional dysregulation.

## Introduction

Sleep alteration is intimately linked to stress-related neuropsychiatric disorders and is increasingly recognized as a mechanistically relevant transdiagnostic process, owing to its shared neurobiological architecture with affective pathology and the emerging therapeutic potential of sleep modulation for affective dysfunction^1–9^. As a stage highly sensitive to stress, REM sleep has been thought to engage neural mechanisms that substantially overlap with those governing threat processing and defensive behavioral responses^10–13^. While these responses are adaptive under acute threat, persistent activation of threat- and stress-responsive systems by chronic stress can become maladaptive, giving rise to anxiety-, fear-, and depression-like affective states^14–16^. Therefore, chronic stress-induced REM sleep dysregulation may represent a shared biological mechanism underlying core threat-related symptoms across multiple psychiatric conditions. However, the neural and molecular pathways through which REM sleep contributes to maladaptive affective states remain poorly understood, limiting efforts to develop mechanism-based therapeutic strategies targeting the core pathology of stress-related disorders.

The immune system constitutes a critical interface linking internal and external challenges to neural regulation, with peripheral cytokines and circulating immune cells modulating brain circuits that govern complex, state-dependent behaviors^17–28^. Within the central nervous system, microglia play a pivotal role in neuroimmune communication by integrating peripheral immune signals and directly regulating neuronal excitability and circuit dynamics^18,20,29,30^. Under chronic stress conditions, accumulating evidence suggests that infiltrating monocytes, extracellular matrix remodeling, and microglial activation contribute to anxiety- and depression-like behavioral states^22,23,31–35^. However, how chronic stress-induced immune activation interacts with region- and cell-type-specific neuronal mechanisms to reshape REM sleep regulation and drive individual differences in stress susceptibility remains poorly understood.

Here we identify stress-induced persistent REM sleep alterations as a state-dependent physiological window associated with affective susceptibility and uncover a neuroimmune feedforward mechanism that perpetuates this stress-associated state. We show that splenic nerve activation promotes monocyte recruitment through the choroid plexus, leading to microglial remodeling and IL6– gp130-mediated potentiation of GPe PV⁺ neuron excitability. State-specific manipulations indicate that GPe PV⁺ neurons during REM sleep influence threat-related behavioral abnormalities. These findings reveal a REM-linked neuroimmune circuit through which peripheral immune activation promotes affective dysfunction.

## Results

### Elevated REM sleep predicts stress susceptibility

To characterize the relationship between increases in REM sleep and stress susceptibility following chronic stress and to elucidate the underlying mechanisms, we employed a chronic predator stress (CPS) model that exposes mice to ethologically relevant stressors (Figure 1A)^10,36^. On day 1 of CPS, NREM sleep was significantly increased relative to baseline, consistent with previous reports^37^. With continued predator exposure, total REM sleep time progressively increased, reaching statistical significance by day 4 of CPS and remaining elevated through day 12. These alterations were accompanied by a concomitant reduction in wakefulness. Substantial inter-individual variability in REM sleep enabled classification of mice into REM sleep-increased (RI) and REM sleep-normal (RN) groups, with RN mice resembling unstressed controls (CON) (Figure 1B and Extended Data Fig. 1A-1C). Stress susceptibility was assessed and correlated with individual REM sleep duration after 12 days of CPS exposure. Threat-related behaviors, such as heightened anxiety and exaggerated stimulus-evoked defensive responses, were used as indicators of behavioral susceptibility to stress. Increased REM sleep was associated with heightened anxiety-like behavior, as indicated by reduced time spent in the center of the open field test (OFT), and with enhanced defensive responses to looming stimuli, characterized by decreased response latency (Figure 1C). Consistent with our previous findings that enhanced threat-related neuronal activity is associated with increased anhedonia, but not social avoidance, increased REM sleep was also associated with reduced sucrose preference (Figure 1D), while showing no significant association with social avoidance behavior (Figure 1D)^38^. These findings suggest that chronic stress-induced REM sleep elevation preferentially predicts stress susceptibility linked to threat-related phenotypes, while remaining dissociable from social deficits^22^. Together, this establishes REM sleep dysregulation as a tractable framework for investigating the mechanisms underlying phenotype-specific stress vulnerability.

**Fig 1.**
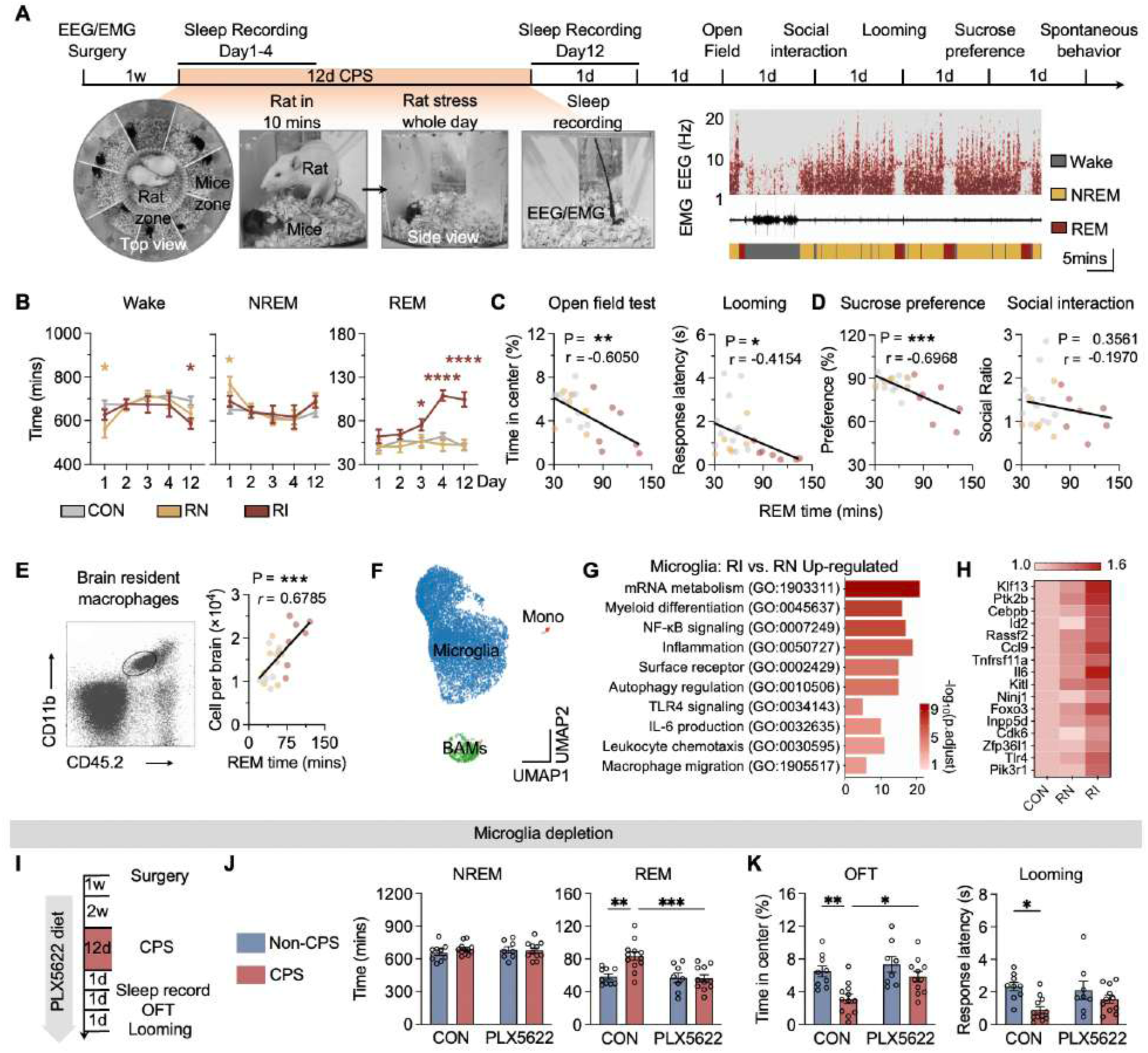
Microglia link REM sleep to stress-induced threat-related behavior. (A) Experimental timeline and schematic of the chronic predator stress (CPS) paradigm. Mice underwent EEG/EMG implantation, followed by 12 days of CPS with sleep recordings on days 1–4 and day 12, and subsequent behavioral tests. (B) Time spent in wakefulness, REM sleep, and NREM sleep across days 1–4 and day 12 of CPS. RI mice (REM sleep-increased) were defined by total REM duration exceeding the control mean + 2 SD threshold; remaining mice were classified as RN (REM sleep-normal). (C) Correlations between total REM sleep duration and threat-related behaviors on day 12 after CPS, including open-field center time and looming response latency. Pearson’s correlation coefficients (r) and p values are indicated. (D) Correlations between total REM sleep duration, sucrose preference, and social interaction ratio on day 12 after CPS. Pearson’s correlation coefficients (r) and p values are indicated. (E) Flow cytometry gating strategy and correlation between the number of brain-resident macrophages per brain and total REM sleep time. (F) UMAP representation of CD11b⁺ myeloid cells showing clusters corresponding to microglia, BAMs, and monocytes. (G) Gene Ontology (GO) biological processes significantly upregulated in microglia from RI mice compared to RN mice. (H) Heat map showing expression of representative differentially expressed genes within the “leukocyte differentiation” pathway across CON, RN, and RI groups (adjusted p < 0.05). (I) Experimental design for microglial depletion using PLX5622 diet during CPS. (J) Time spent in NREM and REM sleep in CON, PLX5622, CON + CPS, and PLX5622 + CPS groups. (K) Open-field (left) and looming (right) behavioral measurements in the indicated groups. All data are shown as mean ± SEM. Statistical significance is indicated as P < 0.05 (*), < 0.01 (**), < 0.001 (***), and < 0.0001 (****). See Table S1.

### GPe microglia modulate stress-induced REM sleep

Given the critical role of neuroimmune interactions, particularly the regulation of stress-related behaviors by microglia and monocyte^20,22,28,39^, we examined their contribution on day 4 of CPS, when RI and RN mice first began to clearly diverge. Flow cytometric analysis revealed a positive association between the abundance of brain-resident macrophages, including microglia, and individual REM sleep duration (Figure 1E). To further characterize the immune landscape involved in stress-induced REM sleep, we performed single-cell RNA sequencing (scRNA-seq) of brain CD11b⁺ cells isolated from RI, RN, and CON mice. Subcluster analysis identified monocyte, microglial, and border-associated macrophages (BAM) populations (Figure 1F). Notably, microglia exhibited a substantially larger set of upregulated genes compared with BAMs (Extended Data Fig. 2A-2B). Gene Ontology analysis of microglial upregulated genes in RI mice revealed enrichment in pathways related to innate immune responses, chemotaxis, and interleukin-6 (IL6) production (Figure 1G and Extended Data Fig. 2C), suggesting activation of inflammatory programs.

**Fig 2.**
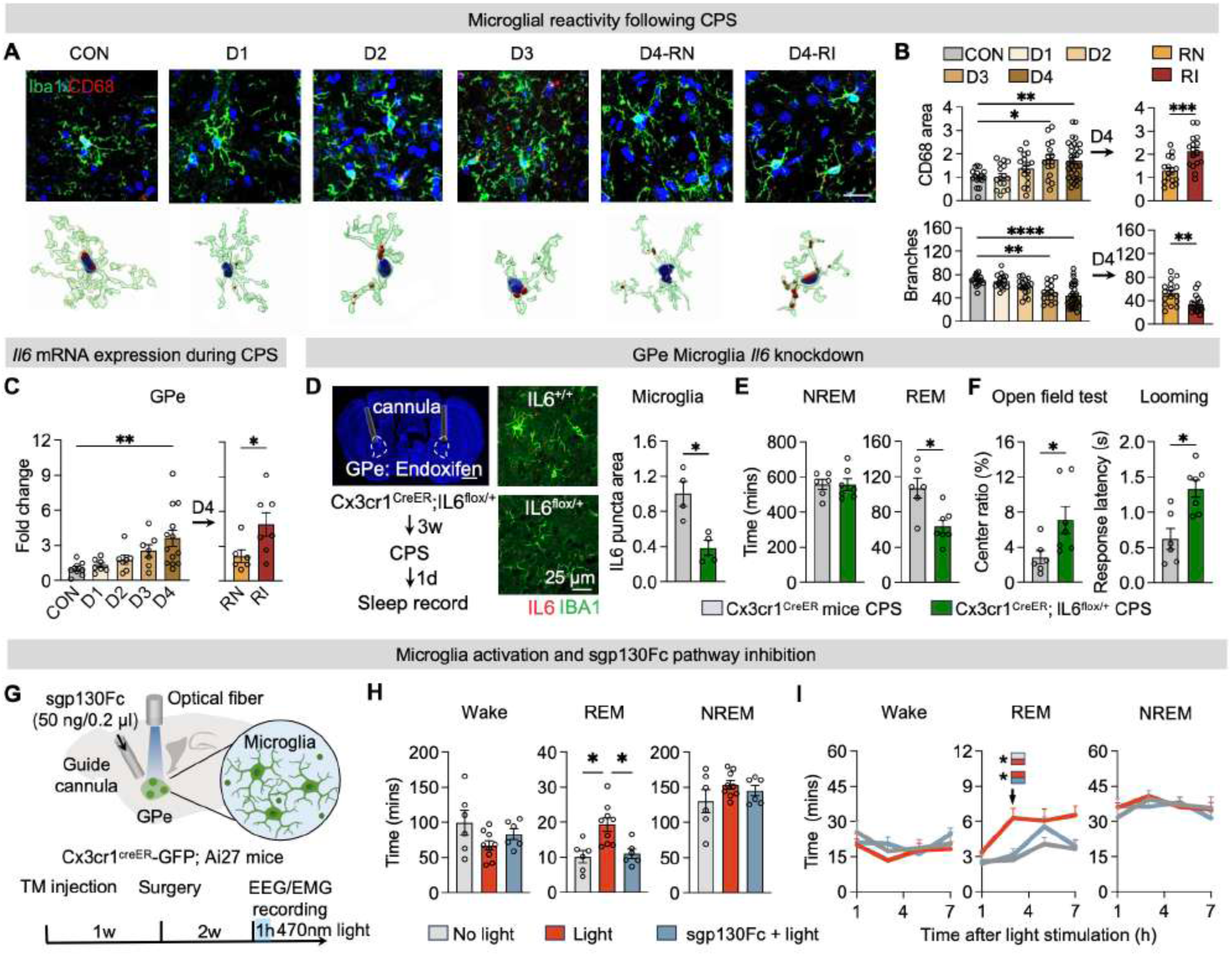
GPe microglia-derived IL6 regulates sleep and threat-related behavioral changes. (A) Representative images of immunostaining and 3D reconstruction of microglial morphology in the GPe from CON mice and CPS for 1–4 days, including RN and RI mice at day 4 (scale bar, 20 μm). (B) Quantification of CD68 volume (normalized to CON) and microglial branch number across CPS. (C) Relative *Il6* mRNA expression in the GPe. (D) Strategy for microglia-specific *Il6* knockdown in the GPe and validation of knockdown efficiency in Cx3cr1^CreER; Il6^flox/+ mice. Left, cannula placement (scale bar, 1 mm); right, IL6 puncta area in Iba1⁺ cells (scale bar, 25 μm), normalized to CON. (E) Time spent in NREM and REM sleep following CPS in control and GPe microglia-specific *Il6* knockdown mice. (F) Open-field (left) and looming (right) behavioral measurements in the indicated groups. (G) Schematic of GPe microglia-specific optogenetic activation and local sgp130Fc infusion in the GPe. Vehicle or sgp130Fc was locally infused 30 min before 470-nm light stimulation. (H) Effects on wakefulness, REM, and NREM sleep under No light, Light, and sgp130Fc + Light. (I) Time course of sleep-state changes after microglial activation. All data are presented as mean ± SEM. Statistical significance is indicated as P < 0.05 (*), < 0.01 (**), < 0.001 (***), and < 0.0001 (****). See Table S1.

**Fig 3.**
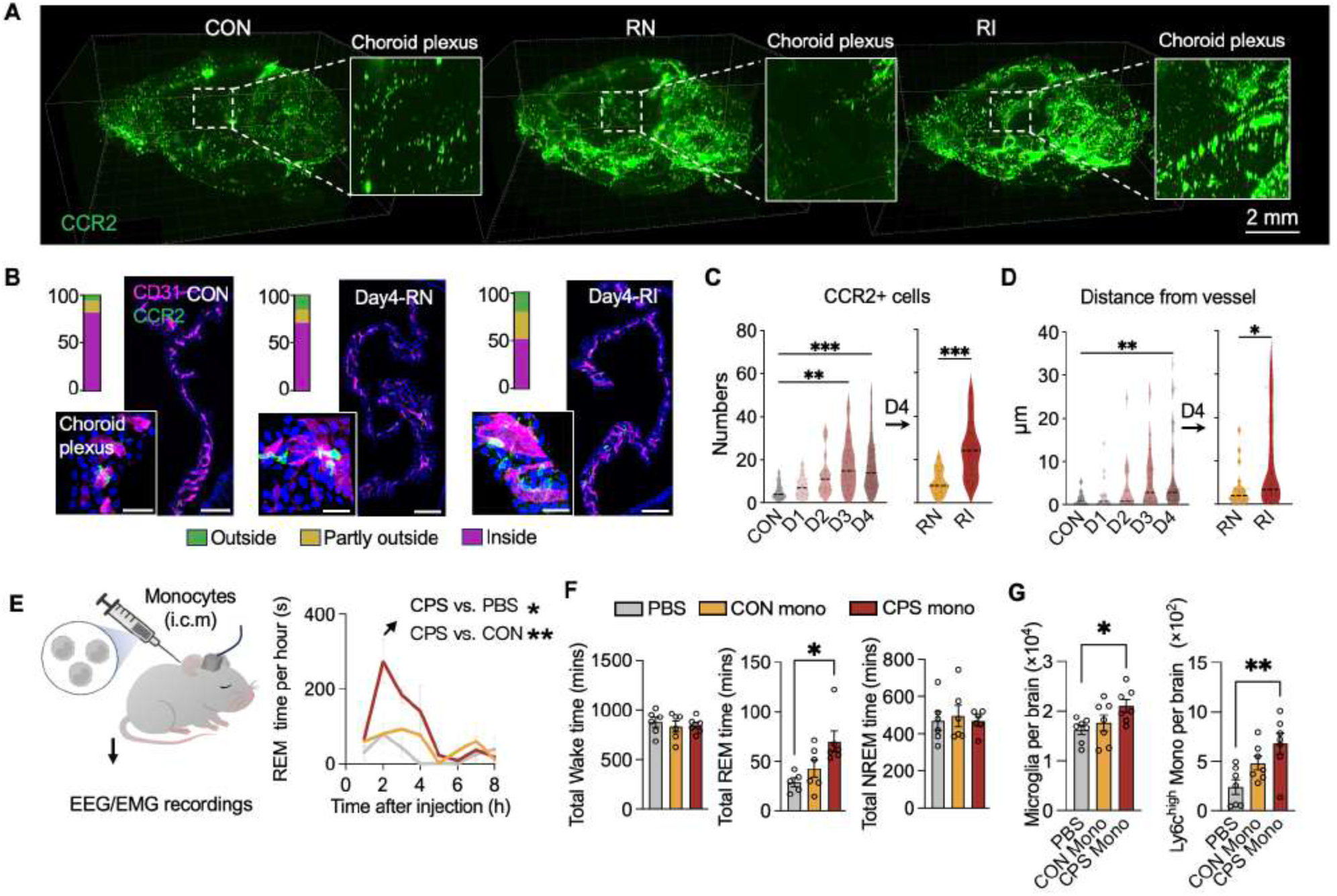
Peripheral monocytes engage a feedforward neuroimmune circuit via brain infiltration following stress. (A) Whole-brain clearing and 3D reconstruction of CCR2⁺ cells in CCR2^GFP-DTR mice across CON, RN, and RI groups (scale bar, 2 mm). (B) Immunofluorescence images showing the distribution of CCR2⁺ monocytes relative to CD31⁺ vasculature (scale bar, 100 μm). Insets show higher-magnification views (scale bar, 20 μm). (C–D) Quantification of CCR2⁺ cell number (C) and distance from blood vessels (D) in CON, RN, and RI mice. (E) Experimental schematic illustrating intracisternal magna (i.c.m.) injection of monocytes followed by EEG/EMG recording, and the time course of REM sleep changes after injection. REM sleep time per hour is shown for mice receiving PBS, monocytes from control donors (CON mono), or monocytes from CPS-exposed donors (CPS mono). (F) Total time spent in wakefulness, REM sleep, and NREM sleep during an 8-h recording following i.c.m. injection of PBS, CON monocytes, or CPS monocytes. (G) Quantification of microglial and Ly6C^high monocyte numbers in the brain after i.c.m. injection of PBS, CON monocytes, or CPS monocytes, assessed by flow cytometry. All data are presented as mean ± SEM. Statistical significance is indicated as P < 0.05 (*), < 0.01 (**), < 0.001 (***), and < 0.0001 (****). See Table S1.

To investigate the role of microglia in stress-induced REM sleep alterations, microglia were pharmacologically depleted using PLX5622, a selective colony-stimulating factor 1 receptor (CSF1R) antagonist^40^. PLX5622 treatment effectively reduced Iba1⁺ cells across the brain (Extended Data Fig. 3). Microglial depletion attenuated CPS-induced REM sleep elevation and mitigated threat-related behavioral phenotypes, without affecting baseline sleep or NREM sleep duration (Figure 1I-1K). Although prior studies reported minimal stress-induced changes in microglial homeostatic or inflammatory gene expression,^22^ these differences likely reflect distinct mechanisms underlying social withdrawal versus REM sleep-associated threat-related behaviors. Consistent with this view, stress-induced anxiety-like behavior is associated with robust microglial activation^23,33^.

Given the distributed neural pathways underlying stress-induced anxiety and REM sleep modulation^39^, we next assessed the regional specificity of microglial activation. Whole-brain quantitative mapping of Iba1⁺ cells revealed preferential microglial engagement in RI mice compared with RN and CON mice, with the most pronounced increase observed in the GPe (Extended Data Fig. 4A and 4B). Notably, the GPe is closely linked to CPS-induced REM sleep alterations and enhanced defensive responses^10^. Within the GPe, CPS gradually increased both total microglial density and CD68⁺ phagolysosomal microglia from day 1 to day 3, suggesting a time-dependent transformation of microglia toward an inflammatory state. Most importantly, this activation predominantly distinguished RI mice from RN and control mice on day 4 (Figure 2A and 2B). Given the selective upregulation of IL6 in microglial clusters from RI mice (Figure 1G), we next tested whether microglial IL6 modulates CPS-induced behavioral changes. Notably, although circulating IL6 protein levels were elevated by stress, they did not differ between RI and RN mice (Extended Data Fig. 5A). In contrast, *Il6* mRNA expression within the GPe was selectively increased in RI mice (Figure 2C). Local intracerebral delivery of recombinant IL6 using an osmotic pump into the GPe was sufficient to promote REM sleep without affecting wakefulness or NREM sleep (Extended Data Fig. 5B and 5C). To interrogate the functional role of GPe microglia in REM sleep regulation, we administered endoxifen into the GPe of Cx3cr1-CreER; Il6^flox/+^, in which endoxifen induction drives microglia-specific IL6 knockdown in the GPe^41^. Local Il6 knockdown reduced IL6 puncta in Iba1⁺ cells and attenuated stress-induced REM sleep increase, accompanied by reductions in threat-related behavioral phenotypes (Figure 2D-2F).

**Fig 4.**
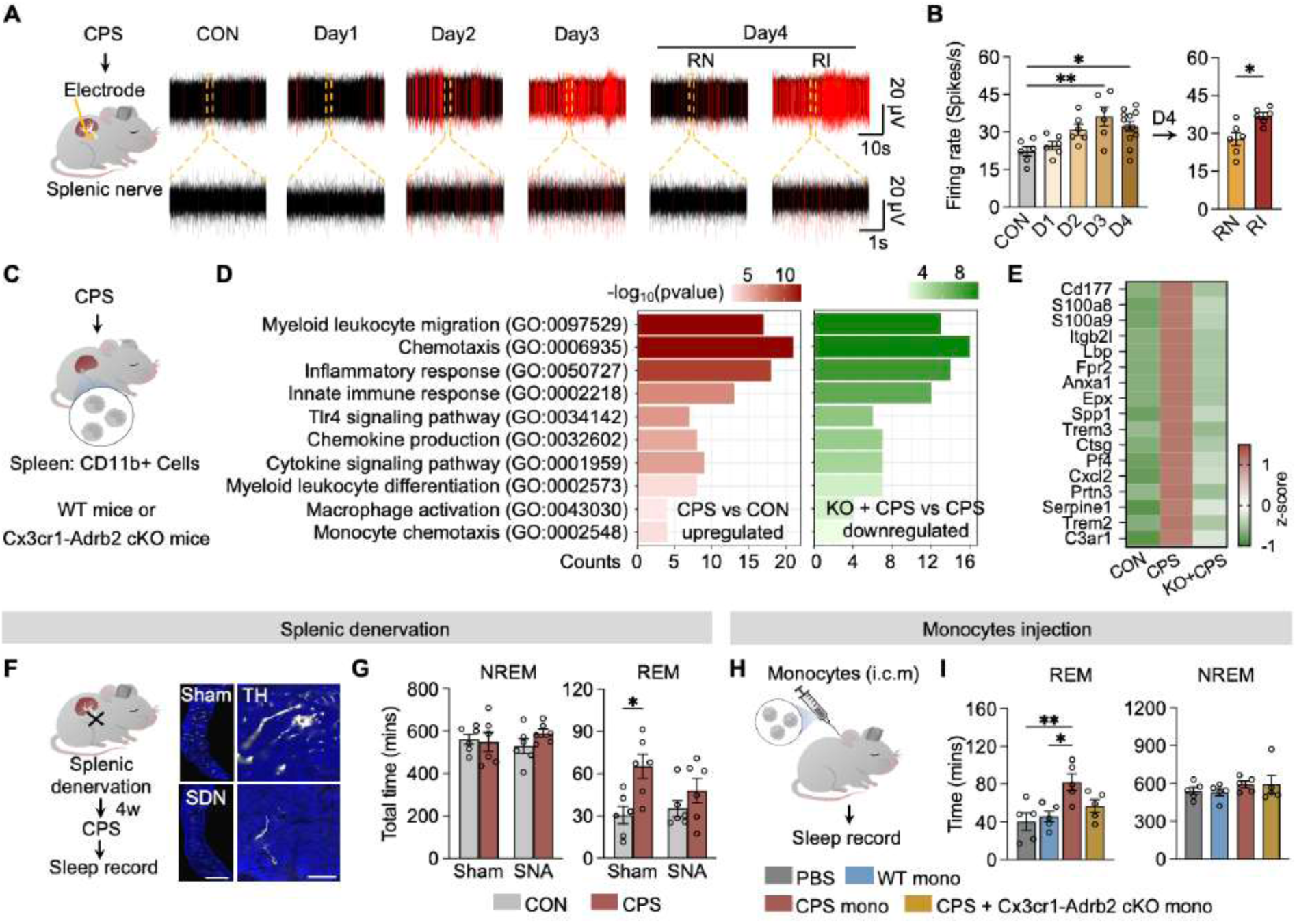
Stress-induced splenic monocyte reprogramming promotes REM sleep augmentation. (A) In vivo recordings of splenic nerve activity during CPS. Spikes (>20 μV) are highlighted. (B) Quantification of splenic nerve firing rate across CON and CPS. (C) Experimental design for isolation of splenic CD11b⁺ cells from CON and from WT or Cx3cr1-Adrb2 conditional knockout mice following CPS. (D) GO enrichment of altered pathways in splenic CD11b⁺ cells (CPS vs CON; KO + CPS vs WT + CPS). CPS-upregulated pathways are shown in red, and pathways reduced upon Adrb2 deletion in green. Nominal p values (uncorrected) are reported for exploratory analysis. (E) Heat map showing Z-scored expression of genes associated with myeloid leukocyte migration (GO:0097529) across CON, CPS, and Cx3cr1-Adrb2 conditional knockout + CPS. (F) Splenic denervation paradigm followed by CPS and sleep recording; Representative tyrosine hydroxylase (TH) staining confirms denervation (scale bars: left, 1 mm; right, 100 μm). (G) Quantification of 24-h total NREM and REM sleep time after splenic denervation under CPS. (H) Experimental schematic of intra-cisterna magna (i.c.m.) injection of circulating monocytes. (I) Effects of i.c.m. injection of CPS-derived monocytes, with or without β2-adrenergic receptor deletion (Cx3cr1-Adrb2 conditional knockout), on total wakefulness, REM sleep, and NREM sleep. All data are presented as mean ± SEM. Statistical significance is denoted as P < 0.05 (*), < 0.01 (**). See Table S1.

**Fig 5.**
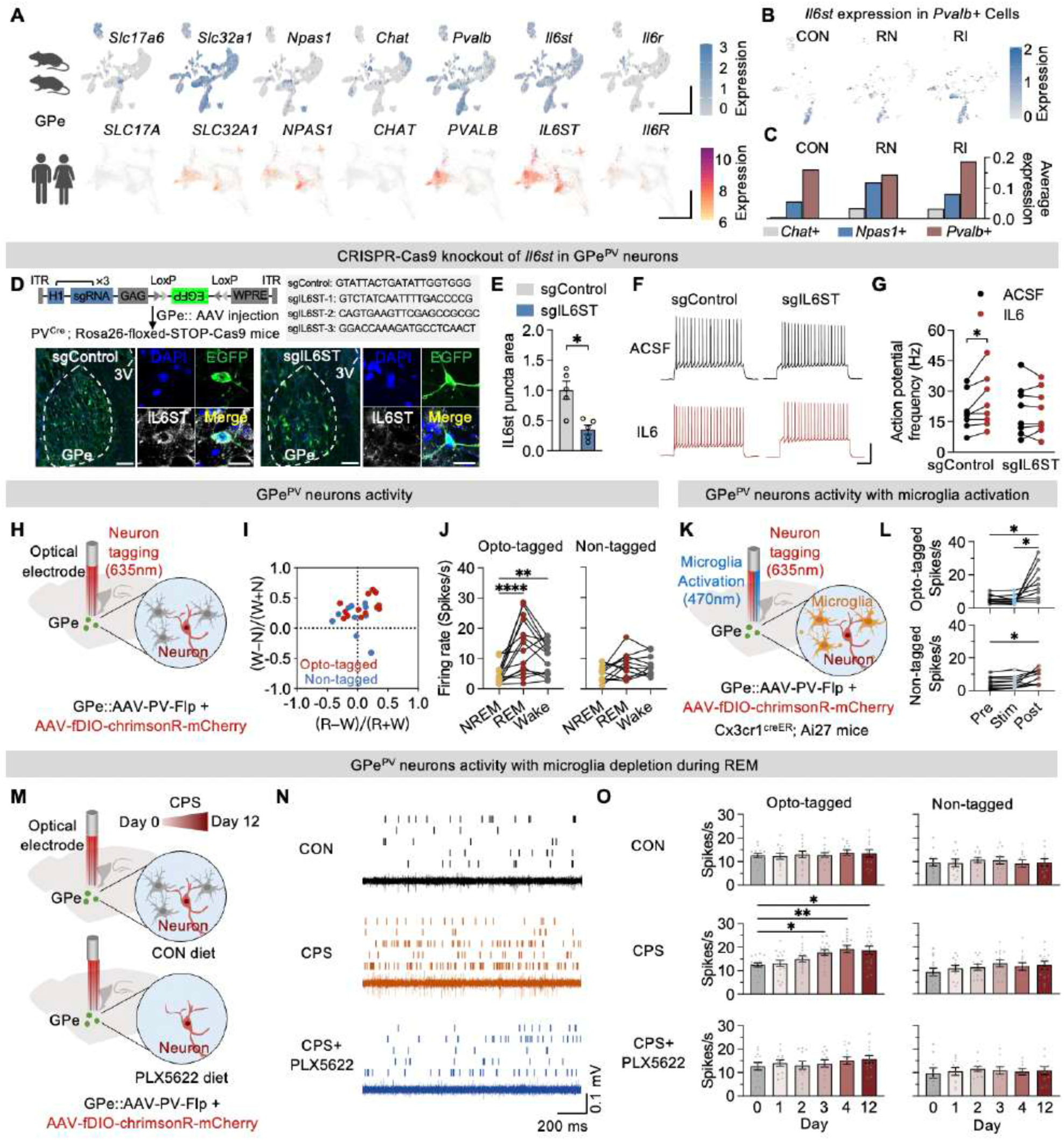
Chronic stress induces sustained activation of GPe PV⁺ neurons through microglia-derived IL-6 signaling. (A) Single-nucleus RNA sequencing UMAP of GPe neurons showing major neuronal populations in GPe (*Pv*⁺, *Npas1*⁺, and *Chat*⁺) and feature plots for Il6st and Il6r, together with canonical glutamatergic (Slc17a6) and GABAergic (Slc32a1) markers in mice (top). Bottom, UMAPs from the Human Brain Atlas showing gene expression in human GPe GABAergic neuron subtypes (SOX6⁺ CTXND1⁺ and LHX6⁺ LHX8⁺ neurons) ^51^. (B) UMAP feature plots showing Il6st expression in mice GPe Pvalb⁺ neurons across CON, RN, and RI groups. (C) Average Il6st expression levels in *Chat*⁺, *Npas1*⁺, and *Pvalb*⁺ GPe neurons across CON, RN, and RI groups. (D) Schematic of viral CRISPR-Cas9-mediated disruption of Il6st using sgIL6ST or non-targeting control sgRNA (sgControl) in PV-Cre; Rosa26-LSL-Cas9 mice (top). Representative immunofluorescence images showing IL6ST puncta and viral EGFP expression in the GPe under sgControl and sgIL6ST conditions (left, scale bar = 200 μm; right, higher magnification, scale bar = 20 μm). (E) Quantification of IL6ST puncta area in the GPe, normalized to sgControl. (F) Representative membrane potential traces from whole-cell recordings of GPe PV neurons under ACSF or IL6 incubation conditions in sgControl and sgIL6ST mice (scale bars: 200 ms, 50 mV). (G) Action potential firing frequency in response to current injections (200 pA, 1000 ms) following ACSF or IL6 incubation. sgControl: n = 8 neurons from 5 mice; sgIL6ST: n = 8 neurons from 6 mice. (H) Schematic of in vivo opto-electrophysiological recording of GPe PV⁺ neurons. (I) Classification of recorded GPe neurons based on state-dependent activity profiles. Scatter plot shows REM–wake versus wake–NREM activity differences (Z-scored firing rates), identifying opto-tagged (red) and non-tagged neurons (blue). Opto-tagged, n = 14 neurons from 4 mice; non-tagged, n = 12 neurons from 4 mice. (J) Firing rates of opto-tagged and non-tagged neurons during NREM, REM, and wakefulness. Lines connect neurons recorded across states. (K) Experimental workflow for microglial optical activation during wakefulness followed by post hoc opto-tagging to classify recorded neurons. (L) Quantification of neuronal firing rates before, during, and after microglial stimulation. (M) Experimental timeline for in vivo electrophysiological recordings during CPS, with or without microglial depletion by PLX5622. (N) Representative spike raster plots and corresponding traces of opto-tagged GPe neurons recorded during REM sleep following 12 days of CPS. (O) Quantification of firing rates of opto-tagged and non-tagged neurons across experimental groups and time points (CON, CPS, CPS + PLX5622; days 0–4 and day 12). All data are presented as mean ± SEM. Statistical significance is denoted as P < 0.05 (*), < 0.01 (**). See Table S1.

To investigate the functional role of GPe microglia in regulating REM sleep, we employed optogenetic stimulation in double-transgenic *Cx3cr1*-CreER-GFP::Ai27 mice, in which tamoxifen induction drives microglia-specific expression of Channelrhodopsin-2 (ChR2)-tdTomato (Figure 2G). Following tamoxifen administration, lineage specificity was confirmed, with ∼95% of GFP⁺ cells co-expressing tdTomato (Extended Data Fig. 5D and 5E). Microglia were optogenetically stimulated using 10-Hz blue light delivered to the GPe, whereas control mice received fiber implantation without light stimulation^42^. Optogenetic stimulation induced pronounced microglial morphological remodeling, characterized by enlarged somata and retracted, less-ramified processes. Quantitative analyses revealed a significant reduction in branch number compared with controls, accompanied by marked upregulation of CD68 expression (Extended Data Fig. 5F and 5G), consistent with a pro-inflammatory shift observed during the CPS. Functionally, although optogenetic stimulation did not elicit an immediate sleep transition, it significantly increased total REM sleep within 4 hours post-stimulation and prolonged the duration of REM episodes, indicating that GPe microglia modulate neuronal circuits governing REM sleep induction in a phase-delayed manner (Figure 2H, 2I, Extended Data Fig. 5H and 5I).

Single-nucleus RNA sequencing (snRNA-seq) revealed minimal expression of the membrane-bound IL6 receptor (*Il6r*) in GPe neurons, whereas the signal-transducing subunit *Il6st* (gp130) was robustly expressed (Figure 5A)^43^. Consistently, local blockade of gp130 signaling in the GPe using sgp130Fc abolished the REM sleep-promoting effects of microglial optogenetic stimulation (Figure 2H and 2I)^43^. Collectively, these findings suggest that IL6 induction reflects a generalized stress response^44–47^, whereas GPe microglia-derived IL6 acts locally to regulate stress-induced REM sleep.

### Neuroimmune feedforward regulation of microglial function by circulating monocytes

Given that stress engages peripheral monocytes to modulate behavior^22,28^, we next examined how they interact with central immune-neural mechanisms underlying REM sleep. Flow cytometric analysis revealed a two-fold elevation of Ly6C^hi^ inflammatory monocytes in both the circulation and brain of RI mice relative to RN and CON mice (Extended Data Fig. 6A-6D). Together, these findings point to a potential contribution of peripheral monocytes to stress-evoked REM sleep and associated behavioral alterations. To test this, we injected Ccr2^GFP-DTR^ mice with diphtheria toxin to deplete Ly6C^hi^ monocytes which are characterized by high CCR2 expression, followed by exposure to CPS. Likewise, the stress-induced elevation in REM sleep was abolished in these mice, while baseline sleep under non-stress conditions remained unchanged. Additionally, we administered a CCR2 antagonist directly into the brain via the cisterna magna during CPS exposure^18^. Brain-specific blockade of CCR2 signaling substantially reduced the infiltration of Ly6C^hi^ monocytes into the brain and mitigated the CPS-induced elevation in REM sleep (Extended Data Fig. 6E and 6F). Interestingly, the CPS-induced increase in microglial numbers was abolished by CCR2⁺ cell depletion or CCR2 blockade (Extended Data Fig. 6F), suggesting that inflammatory monocytes modulate stress-associated microglial activation.

**Fig 6.**
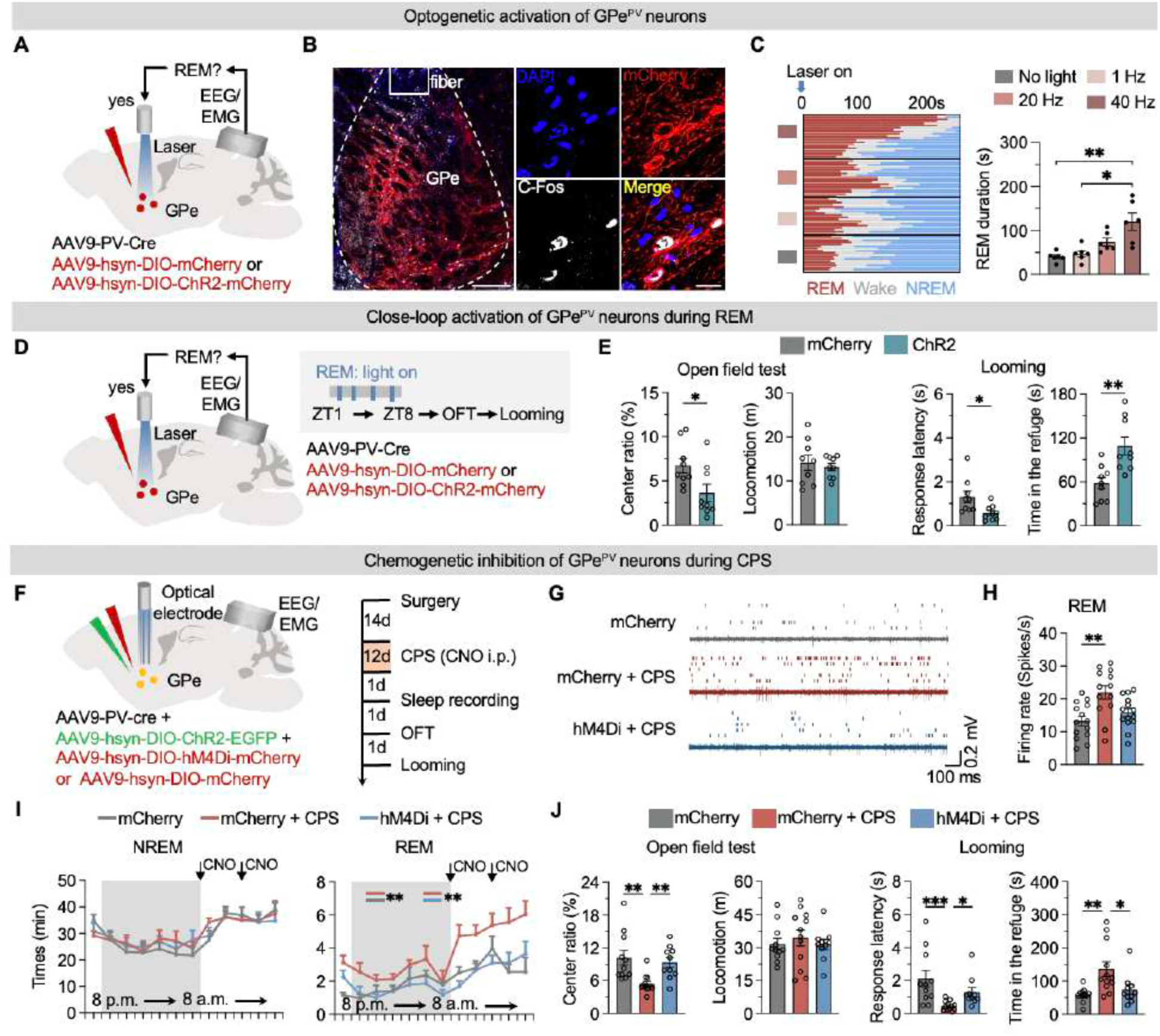
REM-active GPe PV⁺ neurons are required for stress-induced REM sleep and threat-related behaviors. (A) Schematic of the closed-loop optogenetic stimulation paradigm targeting GPe PV neurons selectively during REM sleep, combined with EEG/EMG recording. (B) Representative images showing co-localization of c-Fos and mCherry in GPe neurons following REM-dependent closed-loop optogenetic stimulation (Left, scale bar = 200 μm; Right, scale bar = 20 μm). (C) Left, vigilance-state transitions aligned to laser onset during optogenetic stimulation at different frequencies (no light, 1 Hz, 20 Hz, and 40 Hz). Right, quantification of REM sleep duration across conditions. (D) Schematic of the closed-loop optogenetic stimulation paradigm targeting GPe^PV^ neurons selectively during REM sleep, combined with EEG/EMG recording and subsequent behavioral testing. (E) Effects of REM-specific optogenetic activation of GPe^PV^ neurons on open-field test performance and looming-evoked defensive behavior. (F) Schematic of chemogenetic inhibition of GPe PV⁺ neurons. (G) Representative spike raster plots showing firing activity of GPe PV+ neurons in mCherry control, mCherry + CPS, and hM4Di + CPS conditions. (H) Quantification of REM-state firing rates of GPe PV+ neurons across experimental groups. (I)Time-course of REM sleep (left) and NREM sleep (right) across the light–dark cycle following CPS. Gray shading indicates dark phase. (J) Behavioral performance in the open-field test and looming assay in mCherry, mCherry + CPS, and hM4Di + CPS groups. All data are presented as mean ± SEM. Statistical significance is denoted as P < 0.05 (*), < 0.01 (**) and < 0.001 (***). See Table S1.

We next used multimodal imaging in Ccr2^GFP-DTR^ mice to map Ly6C^hi^ inflammatory monocyte trafficking in the brains of RI, RN, and CON mice on day 4 of CPS. We began with whole-brain tissue clearing using the iDISCO method, followed by three-dimensional light-sheet microscopy of brains to detect GFP⁺ monocytes. Whole-brain analysis revealed a pronounced accumulation of GFP⁺ cells in the choroid plexus (ChP) lining of RI mice, accompanied by infiltration into the ventricular system (Figure 3A and Extended Data Fig. 7B). Quantitative analysis confirmed that GFP⁺ cell numbers in the ChP progressively increased from day 1 to day 3 of CPS, with a significant elevation observed on day 4 in RI mice compared with both RN and CON groups. At this time point, a large proportion of monocytes were extravascular, indicating active emigration from the vasculature (Figure 3B-3D and Extended Data Fig. 7A). In contrast, no significant differences in GFP⁺ cell abundance were detected in the meninges between RI and RN mice (Extended Data Fig. 7B-7E).

To determine whether circulating monocytes intrinsically acquire brain-infiltrating capacity under stress, we performed adoptive transfer experiments. Ccr2^GFP-DTR^ mice were subjected to CPS and stratified by REM sleep phenotype as RI-GFP or RN-GFP. CD11b⁺ cells were isolated from the peripheral blood of these donor mice and intravenously transferred into naïve wild-type recipients as well as CPS-exposed RI and RN mice (Extended Data Fig. 8A). RI-GFP⁺ donor cells accumulated significantly more in the ChP of recipient mice than RN-GFP⁺ donor cells. Notably, RI-GFP⁺ cells accumulated to a similar extent in the ChP of both RI and RN recipients (Extended Data Fig. 8B and 8C). These findings indicate that chronic stress primes the brain for peripheral monocyte recruitment, whereas the intrinsic inflammatory and chemotactic properties of circulating monocytes determine the REM sleep elevation phenotype.

To further test the functional significance of monocyte entry into the brain, we isolated monocytes from the peripheral blood of CON and CPS-exposed mice and introduced them into the cerebrospinal fluid (CSF) of naïve WT recipients via intracisternal injection (Figure 3E). Injection of WT monocytes produced a modest increase in REM sleep, whereas delivery of CPS-derived monocytes induced a robust, approximately two-fold increase in REM sleep that persisted for up to 4 hours (Figure 3F). Moreover, two hours after injection, mice receiving CPS monocytes displayed a markedly greater number of Ly6C^hi^ monocytes in the brain, accompanied by increased microglial numbers, compared to controls (Figure 3G), suggesting infiltrating monocytes act as upstream triggers of stress-induced microglial activation.

We next examined how stress signals are conveyed to circulating monocytes. The spleen serves as a major reservoir for inflammatory monocytes mobilized during psychological stress and implicated in behavioral regulation^24,33,48,49^. Electrophysiological recordings revealed a gradual increase in splenic nerve activity as CPS progressed from day 1 to day 4, with activity peaking at day 3, preceding both the peak of monocyte-associated inflammatory signaling in the brain and the subsequent emergence of REM sleep elevation. Notably, on day 4, splenic nerve activity remained significantly elevated in RI mice compared with RN and CON mice (Figure 4A-4B), suggesting sustained sympathetic activation in RI mice. Consistent with this, splenic denervation attenuated stress-induced

REM sleep without affecting baseline REM sleep (Figure 4F-4G). To define the molecular consequences of stress-evoked sympathetic signaling in splenic monocytes, we performed bulk RNA sequencing of CD11b⁺ splenic cells from WT and Cx3cr1-Adrb2 cKO mice, which lack β2-adrenergic receptor signaling in myeloid cells. GO analysis showed that CPS robustly induced inflammatory and chemotactic gene programs in WT splenic CD11b⁺ cells, whereas this induction was markedly attenuated in CD11b⁺ splenic cells from Cx3cr1-Adrb2 cKO mice (Figure 4D-4E). Consistent with this, intracisternal injection of circulating CD11b⁺ cells from CPS-exposed Cx3cr1-Adrb2 cKO mice into naïve WT recipients failed to elicit REM sleep elevation compared with cells from CPS-exposed WT donors (Figure 4H-4I), demonstrating that β2-adrenergic signaling is required for the acquisition of this sleep-modulating phenotype. Together, these findings support a role for splenic neural inputs in priming inflammatory monocytes that contribute to stress-induced alterations in REM sleep.

### Microglia sustain GPe PV⁺ neuron activity following CPS

To determine how GPe neurons respond to microglia-derived IL6, we first analyzed both mouse and human single-cell transcriptomic datasets. The GPe is primarily composed of three major neuronal populations, including Pvalb⁺, Npas1⁺, and Chat⁺ neurons^50^. In our mouse GPe snRNA-seq dataset, *Il6st* expression was preferentially enriched in Pvalb⁺ neurons compared with Npas1⁺ and Chat⁺ populations (Figure 5A-5C). Notably, *Il6st* expression was further elevated in RI mice relative to RN and CON groups, with the strongest increase in the Pvalb⁺ population (Figure 5B-5C), consistent with a state-dependent enhancement of IL6 signaling. Consistently, analysis of the Human Brain Atlas^51^ revealed conserved enrichment of IL6ST expression in human GPe PVALB+ neuronal populations, suggesting cross-species conservation of IL6 sensitivity in GPe neurons (Figure 5A).

We then investigated whether GPe PV⁺ neuron excitability is mediated through the IL6/IL6ST signaling pathway, by achieving PV⁺ neuron-specific CRISPR-Cas9-mediated knockdown of IL6ST. PV-Cre mice were crossed with Rosa26-LSL-Cas9 mice, and AAVs encoding either Il6st-targeting guide RNAs (sgIL6ST) or control guide RNAs (sgControl) were stereotaxically injected into the GPe (Figure 5D). Immunofluorescence confirmed a marked reduction of IL6ST expression in GPe PV⁺ neurons in sgIL6ST-injected mice compared with sgControl-injected mice (Figure 5D and 5E). In vector control neurons, bath application of recombinant IL6 significantly increased the number of current-evoked action potentials in GPe PV⁺ neurons, and this effect was abolished by IL6ST knockdown (Figure 5F-5G). Furthermore, IL6ST knockdown in GPe PV⁺ neurons attenuated stress-induced REM sleep elevation and reduced associated threat-related behavioral phenotypes (Extended Data Fig. 9A and 9B). These results suggest a role for GPe PV⁺ neurons in microglia-IL6-dependent modulation following stress.

To examine PV⁺ neuronal activity in vivo, we performed opto-electrophysiological recordings. AAVs encoding PV-Flp and Flp-dependent ChrimsonR were injected into the GPe, intermittent 40-Hz 635-nm light pulses were delivered, and single units with reliable, short-latency, low-jitter light-evoked responses were identified as optotagged PV⁺ neurons (Extended Data Fig. 10B). Simultaneous EEG/EMG recordings revealed that opto-tagged PV⁺ neurons exhibited state-dependent activity, with maximal firing during REM sleep, whereas non-optotagged (non-PV⁺) neurons did not display this pattern (Figure 5I and 5J). At the population level, firing rates of GPe PV⁺ neurons increased consistently during NREM→REM and NREM→wake transitions and decreased during REM→wake and wake→NREM transitions (Extended Data Fig. 10D), indicating tight coupling to REM sleep state. We next examined how microglia modulate GPe PV⁺ neurons response to microglial stimulation.

AAVs encoding PV-Flp and Flp-dependent ChrimsonR were injected into the GPe of Cx3cr1-CreER::Ai27 mice (Figure 5K). This strategy enabled optotagging of PV neurons using 635 nm light during in vivo electrophysiological recordings and optogenetic activation of microglia using 470 nm light. We optogenetically stimulated GPe microglia while recording PV⁺ neuronal activity (Figure 5K-L and Extended Data Fig. 10E). PV⁺ neurons exhibited stronger responses to microglial stimulation than unidentified neurons, with firing rates increasing to 20-30 Hz.

We subsequently assessed the endogenous response of GPe PV⁺ neurons to CPS and evaluated the contribution of microglia by pharmacologically depleting them with PLX5622 (Figure 5M). The endogenous firing rates of PV⁺ neurons increased by day 3, peaked on day 4, and remained elevated through day 12 of CPS (Figure 5N-5O and Extended Data Fig. 10F), closely paralleling the emergence of the REM sleep phenotype. Notably, under this CPS-induced condition, GPe PV⁺ neurons exhibited an average firing rate of ∼20 Hz, with peak activity reaching up to ∼30 Hz, consistent with that observed during microglial optogenetic stimulation (Figure 5L and 5O). Supporting this, in microglia-depleted mice, PV⁺ neuronal activity during REM sleep failed to increase as CPS progressed (Figure 5N-5O and Extended Data Fig. 10F). Together, these findings suggest that stress primes microglia to drive a sustained elevation of GPe PV⁺ neuronal activity, thereby promoting a stress-induced increase in REM sleep.

### GPe PV⁺ neurons mediate stress-induced REM sleep and associated threat-related behaviors

Next, we investigated the functional role of GPe PV⁺ neurons in REM sleep regulation. We injected AAVs encoding PV promoter-driven Cre recombinase and Cre-dependent ChR2-mCherry into the GPe of WT mice (Figure 6A). mCherry-positive cells coexpressed endogenous PV, confirming the specificity and efficiency of the targeting strategy (Extended Data Fig. 11A and 11B). Functional ChR2 expression was validated by cFos immunostaining following laser stimulation (Figure 6B). Unilateral laser stimulation (1, 20, or 40 Hz) was applied during stable NREM or REM sleep. During NREM sleep, optogenetic stimulation of GPe PV⁺ neurons transiently increased wakefulness without inducing REM sleep. However, in a closed-loop protocol triggered at REM onset and terminated at REM offset, 20- and 40-Hz stimulation prolonged REM episode duration (Figure 6C and Extended Data Fig. 11C-11D). Given the dual wake- and REM-associated functions of this neural population, we examined whether their activity during the two states differentially influences behavioral outcomes. Closed-loop stimulation confined to REM sleep during a single light-phase interval heightened anxiety-like behavior in the OFT, and enhanced defensive responses to looming stimuli after awakening (Figure 6D-6E). In contrast, optogenetic activation immediately before behavioral testing during wakefulness did not alter these threat-related behaviors (Extended Data Fig. 11F-11H), demonstrating that the behavioral impact of GPe PV⁺ neurons is selectively gated by REM sleep.

To assess the contribution of elevated GPe PV⁺ neuron activity during CPS, we chemogenetically inhibited this population throughout stress exposure. WT mice received bilateral GPe injections of AAV-DIO-hM4Di-mCherry or control AAV-DIO-mCherry together with AAV-PV-Cre, and were subsequently subjected to CPS or maintained under non-stressed conditions (Figure 6F). CNO-mediated inhibition was validated by reduced firing of GPe PV⁺ neurons. CPS induced a significant increase in GPe PV⁺ neuronal firing in control mice, whereas this increase was markedly attenuated in hM4Di-expressing mice (Figure 6G-6H). Consistent with these findings, the stress-induced increase in total REM sleep time observed in control mice was abolished in hM4Di mice (Figure 6I). hM4Di mice subjected to CPS spent more time in the center of the OFT and exhibited reduced defensive responses to looming stimuli compared with stressed control mice, resembling the behavioral profile of non-stressed controls (Figure 6J). Together, these results suggest that GPe PV⁺ neuron activity during REM sleep is selectively coupled to stress-related behavioral output, positioning REM sleep as a state-dependent physiological substrate through which behavioral vulnerability may be modulated.

## Discussion

Our previous work identified a medial subthalamic nucleus-GPe pathway that coordinates REM sleep and defensive responses during sustained predator stress, facilitating defensive responses to subsequent threats^10^. Here, we identify GPe PV⁺ neurons as a stress-responsive population central to this circuit and show that their recruitment is dynamically primed by immune signals. Stress-evoked splenic neural activity engages peripheral monocytes, which amplify GPe PV⁺ activity via a neuroimmune feedforward loop, enhancing REM sleep and exacerbating anxiety-like and defensive behaviors. Disrupting this immune component through microglial depletion or blockade of monocyte infiltration abolishes stress-induced increases in REM sleep and associated behaviors without affecting baseline sleep architecture. Importantly, state-specific manipulations further indicate that GPe PV⁺ neurons shape affective behavior preferentially during REM sleep, supporting the idea that REM sleep constitutes an actionable brain state in which immune-primed circuits exert behavioral control. Together, these findings define a new mechanistic framework in which neuroimmune interactions govern the coupling between REM sleep regulation and threat-related behaviors.

Under conditions of imminent threat, increased REM sleep, heightened anxiety, and defensive responses mediated by neuroimmune circuits may confer adaptive advantages by facilitating threat anticipation, promoting vigilance, and enhancing risk avoidance, particularly during injury or infection associated with predation, without disrupting overall sleep continuity^10,11^. In contrast, in modern contexts where stress is predominantly chronic, such as persistent psychological stressors or low-grade inflammation, and rarely linked to immediate survival threats (e.g., predation), persistent engagement of these evolutionarily conserved circuits may represent an evolutionary mismatch, leading to maladaptive outcomes and contributing to anxiety-, fear-, and depression-related pathology^11,14^. Within this framework, our results suggest that neuroimmune feedforward circuits bias the system toward sustained activation, thereby promoting stress susceptibility and highlighting immune-targeted approaches as potential therapeutic strategies for stress-related sleep disturbances and neuropsychiatric comorbidities.

### Data and code availability

Single-cell and bulk RNA sequencing data have been deposited at GEO and are publicly available as of the date of publication. This study also utilized Human and Mammalian Brain Atlas: Basal Ganglia^51^. Further information and requests should be directed to and will be fulfilled by lead contact.

## Acknowledgements

We thank Chenyan Ma and Bo Feng for their valuable comments. This work was supported by grants from Brain Science and Brain-like Intelligence Technology-National Science and Technology Major Project (2025ZD0215000), the Shenzhen Medical Research Fund (B2302004), the National Natural Science Foundation of China (32422036 and 32230042), the Financial Support for Outstanding Talents Training Fund in Shenzhen, and the Shenzhen Science and Technology Program (JCYJ20220530154412028), Guangdong Provincial Key Laboratory of Brain Connectome and Behavior (2023B1212060055) and Shenzhen Brain Science Infrastructure.

## Author contributions

Conceptualization and supervision, Y.-T.T.; methodology, B. Z., S.X. and Y.-T.T.; investigation, B. Z., S.X., L.N.W., J.L., X.Z., Y.X. and Q.H.; writing-original draft, Y.-T.T. with feedback from L.W., X.Y. and H.K.; funding acquisition and resources, L.W and Y.-T.T.

## Competing interests

The authors declare no competing interests.

**Correspondence and requests for materials** should be addressed to Yu-Ting Tseng and Liping Wang.

## Methods

## Animals

Animal care and all experimental protocols were approved by the Animal Care and Use Committees at the Shenzhen Institute of Advanced Technology, Chinese Academy of Sciences. C57BL/6J mice and male Sprague-Dawley rats were obtained from Beijing Vital River. Ccr2-DTR-GFP (Cat. No. NM-KI-210102), Il6-KO mice (Cat. No. NM-KO-18029) and Adrb2-Flox mice (Cat. No. NM-CKO-2114375) were purchased from Shanghai Model Organisms Center, Inc. Cx3cr1-CreER mice (Cat. No. JAX021160), PV-Cre (Cat. No. JAX 008069), RCL-hChR2(H134R)/tdTomato mice (Ai27, Cat. No. JAX012567) were obtained from Jackson Laboratory. Rosa26-CAG-LSL-Cas9-tdTomato mice (Cat. No. T002249) were obtained from GemPharmatech Co., Ltd. For microglia-specific conditional deletion experiments, Cx3cr1-CreER mice were crossed with Il6-flox mice to generate Cx3cr1-CreER; Il6^flox/+^ mice, respectively. For microglia-specific optogenetic activation experiments, Cx3cr1-CreER mice were crossed with Ai27 mice to generate Cx3cr1-CreER; Ai27 mice. For CRISPR-Cas9-mediated deletion in PV neurons, PV-Cre mice were crossed with Rosa26-CAG-LSL-Cas9 mice to generate PV-Cre; Rosa26-LSL-Cas9 mice. All the experiments were performed on adult mice (8–16 weeks of age) of both sexes. Mice were housed in 12-h light/dark cycle (lights on at 08:00 and off at 20:00) with free access to food and water.

### Surgery

Surgical procedures were performed as previously described^10^. Briefly, surgeries were carried out under isoflurane anesthesia (4% induction, 1.5% maintenance) using a stereotaxic frame (RWD, China). For CRISPR-Cas9-mediated knockdown and chemogenetic inhibition experiments using hM4Di, animals received bilateral injections of viral mixtures (100 nl per site); all other experiments involved unilateral injections. The stereotaxic coordinates for injections targeting the globus pallidus externus (GPe) were as follows (in mm relative to bregma): anteroposterior (AP) −0.25, mediolateral (ML) 2.0, and dorsoventral (DV) −4.0. For optogenetic manipulation, a 200-μm optic fiber (NA 0.37; NEWDOON, Hangzhou, China) was implanted 0.1 mm above the viral injection site. The entire assembly was secured with dental cement after implantation.

For electroencephalogram (EEG) and electromyogram (EMG) recordings, an EEG/EMG headmount (Pinnacle, catalog #8201) was affixed to the skull using four stainless steel screws. Two screws (Pinnacle, catalog #8209) were inserted 1.5 mm lateral to the midline and 1.0 mm anterior to bregma, and the other two screws (Pinnacle, catalog #8212) were placed 1.5 mm lateral to the midline and 3.0 mm posterior to bregma. Two EMG wire electrodes were inserted into the neck musculature, and the entire assembly was then sealed with dental cement. Mice were allowed to recover for at least three weeks following viral injection, including at least one week after EEG/EMG implantation before behavioral testing or electrophysiological recordings.

### Virus injections

For optogenetic activation experiments, AAV9-PV-Cre (1.5 × 10¹² genome copies (GC)/mL; BrainVTA, Wuhan) was mixed at a 1:1 ratio with either AAV9-hSyn-DIO-ChR2-mCherry (3.2 × 10¹² GC/mL; Taitool Bioscience, Shanghai) or AAV9-hSyn-DIO-mCherry (3.0 × 10¹² GC/mL; Taitool Bioscience, Shanghai) and unilaterally injected into the GPe of wild-type mice. For CRISPR-Cas9-mediated Il6st knockdown, AAV2-H1-sgIl6st.sp(MCS)×3-CAG-DIO-eGFP-WPRE-pA (2.3 × 10¹² GC/mL; Taitool Bioscience, Shanghai) or AAV2-H1-sgControl.sp(MCS)-CAG-DIO-eGFP-WPRE-pA (2.1 × 10¹² GC/mL; Taitool Bioscience, Shanghai) was bilaterally injected into the GPe of PV-Cre; Rosa26-floxed-STOP-Cas9 mice. For optrode recordings with optogenetic tagging, AAV-PV-Flp (1.5 × 10¹² GC/mL; BrainVTA, Shanghai) was mixed 1:1 with AAV-fDIO-ChrimsonR-mCherry (3.6 × 10¹² GC/mL; BrainVTA, Shanghai) and unilaterally injected into the GPe of Cx3cr1-CreER; Ai27 mice or wild-type mice. For chemogenetic inhibition of GPe neurons, AAV9-PV-Cre (1.5 × 10¹² GC/mL; BrainVTA, Wuhan) was mixed 1:1 with either AAV9-hSyn-DIO-hM4Di-mCherry (3.3 × 10¹² GC/mL; Braincase, Wuhan) or AAV9-hSyn-DIO-mCherry (3.0 × 10¹² GC/mL; Taitool Bioscience, Shanghai) and bilaterally injected into the GPe of wild-type mice.

### Cisterna magna injection and cerebrospinal fluid collection

Cisterna magna injection was performed as previously described^18^. Briefly, mice were anesthetized with ketamine (100 mg/kg) and xylazine (10 mg/kg) and placed in a stereotaxic frame to expose the foramen magnum. A ∼1 cm midline incision was made in the scalp, and the underlying musculature was gently separated to expose the cisterna magna. A Hamilton syringe was used to slowly inject 5 μL of saline, anti-CCR2 antibody (Sigma-Aldrich, #227016; 1 μg per injection).

For monocyte transplantation experiments, monocytes were isolated from peripheral blood mononuclear cells using magnetic-activated cell sorting (MACS) negative selection with the MojoSort™ Mouse Monocyte Isolation Kit (BioLegend, #480154). A total of 3 × 10⁴ cells suspended in 5 μL of PBS were injected into the cisterna magna.

Cerebrospinal fluid (CSF) collection was performed using the same surgical approach, except that the syringe was operated in aspiration mode. Approximately 8–10 μL of CSF was collected from each mouse, and only clear, blood-free samples were retained for subsequent analyses. After removal of the syringe, the incision was sutured, and mice were placed on a heating pad for recovery.

### Osmotic pump implantation

Osmotic pump implantation was performed as previously described^43^ to target the GPe. Osmotic pumps (RWD, 1002W; 2-week infusion duration, 0.25 μL/hour) were coupled with a Brain Infusion Kit (RWD, BIC-3) and prepared according to the manufacturer’s instructions. Recombinant mouse IL6 (R&D Systems, 40-6ML-200CF) was dissolved in sterile artificial cerebrospinal fluid (aCSF) at a concentration of 2 ng/mL. Pumps were filled with 100 μL of aCSF or IL6-containing aCSF, and the infusion tubing (2 cm) was backfilled before connection. After scalp incision, a subcutaneous pocket was created between the skin and underlying muscle of the back to accommodate the osmotic pump. The infusion cannula was positioned stereotaxically above the GPe. The pump was gently inserted into the subcutaneous pocket, and the cannula was secured to the skull with dental cement. The skin was repositioned to cover the cemented area without excessive tension.

### Guide cannula implantation and local infusion

For local pharmacological or inducible recombination experiments in the GPe, a guide cannula (RWD, 62004) was implanted bilaterally above the GPe under isoflurane anesthesia using the same stereotaxic coordinates as optic fiber implantation. The cannula was secured to the skull with dental cement. For local induction of CreER-mediated recombination, endoxifen (300 nL, 2 mg/mL) was infused into the GPe of Cx3cr1-CreER; Il6^flox/+^ once daily for 3 consecutive days. ^41^ Mice were allowed to recover for 3 weeks following local endoxifen infusion before subsequent experiments. For blockade of IL6 trans-signaling, sgp130Fc (50 ng in 0.2 μL, R&D systems, 468-MG-100) or ACSF was locally infused into the GPe through the implanted cannula 30 min prior to optogenetic stimulation. ^43^

### Adoptive transfer of monocytes

Peripheral blood mononuclear cells (PBMCs) were isolated from CCR2-DTR-GFP mice using a Mouse Peripheral Blood Lymphocyte Isolation Solution Kit (Solarbio Life Science, P8620). Monocytes were subsequently purified from PBMCs by MACS negative selection using the MojoSort™ Mouse Monocyte Isolation Kit (BioLegend, #480154). Purified monocytes were resuspended in sterile saline, and recipient mice received tail vein injections of 1 × 10^6^ monocytes in 0.2 mL saline.

### EEG/EMG recordings and analysis of sleep-wake states

EEG/EMG recordings were performed as previously described^10^. At least one week after EEG/EMG implantation, behavioral experiments were conducted inside a sound-attenuated chamber using the EEG/EMG recording system (Pinnacle, model 8400-K1). Recordings were initiated after a two-day habituation period. Signals were acquired using a Pinnacle high-gain preamplifier (gain ×100) with a high-pass filter set at 0.5 Hz for EEG and 10 Hz for EMG, and digitized at a sampling rate of 2000 Hz. The raw data were converted into the European data format (.edf). Sleep-wake states were scored using sleep analysis software (SleepSign, Kissei Comtec). EEG and EMG signals were band-pass filtered at 0.25-100 Hz and 30-300 Hz, respectively. The EEG power spectrum for each 4-second epoch was computed by Sleep Sign using the Fast Fourier Transform (FFT) algorithm. Automated sleep staging was performed based on EEG and EMG waveform characteristics within each 4-second epoch. EEG power in the 0.5-4 Hz and 5-8 Hz ranges was defined as delta (δ) and theta (θ) power, respectively. The classification criteria for vigilance states were as follows: wakefulness– presence of movement, high EMG amplitude, and a moderate θ/δ ratio; non-rapid eye movement (NREM) sleep–absence of movement, low EMG activity, and dominance of δ waves; rapid eye movement (REM) sleep–absence of movement, low EMG activity, and dominance of θ waves. All sleep–wake classifications automatically assigned by Sleep Sign were visually inspected and manually corrected when necessary. The total duration and number of episodes for wakefulness, NREM sleep, and REM sleep were exported as text files. Spectrograms were generated using Sirenia Sleep Pro software (Pinnacle).

For different experimental paradigms, recording durations varied. For sleep recording after experiencing chronic predator stress, daily EEG/EMG recordings lasted 23 h and were used for analysis to accommodate the ∼1 h stress procedure. For cisterna magna injection experiments, EEG/EMG data acquired between 12 and 23 h after injection were analyzed; recording durations were kept consistent within each experiment. Total time spent in wakefulness, NREM sleep, and REM sleep, as well as episode numbers, were exported for further analysis. RI mice were defined as those whose total REM duration exceeded the upper physiological range observed in control animals (mean + 2 SD of the control group), whereas the remaining animals were classified as RN.

### Chronic predator stress (CPS)

CPS model was adapted from a previously described paradigm with modifications^10^. Mice were housed in a transparent circular arena (60 cm diameter) containing a central perforated cylindrical compartment (30 cm diameter) that housed two adult male Sprague–Dawley rats, while eight peripheral compartments each contained a single mouse, allowing continuous visual, olfactory, and auditory exposure. Control mice were housed in an identical arena without rats and placed in a separate room. Mice were habituated to the apparatus for 2 days before stress induction. On designated days, rats were introduced into mouse compartments for 10 min to allow close physical contact and then returned to the central compartment, where they remained continuously. Different rats were used each day, and interactions were monitored to prevent injury. For the 4-day CPS protocol, close-contact exposure was performed on days 1–3 but omitted on day 4. For the 12-day CPS protocol, close-contact exposure was omitted on days 4, 8, and 12, with 10-min contact sessions on all other days.

For experiments involving microglial depletion, mice were fed a PLX5622-containing diet or control AIN-76A diet (SyseBio, D20010801) for 2 weeks before the onset of CPS and maintained on the same diet throughout the stress period.

### Behavior tests

Mice were habituated to the testing room for 1 h before behavioral assays. Tests were performed during daylight under dim lighting conditions.

### Open field test

Mice were allowed to freely explore a square open-field arena (50 × 50 × 50 cm) for 5 min. Behavior was recorded using a ceiling-mounted camera providing a top-down view of the entire arena. Locomotor activity and time spent in the center zone (25 × 25 cm) and peripheral zone were quantified using the ANY-maze video tracking system (Stoelting, IL, USA).

### Looming stimulus test

The looming stimulus tests were performed as previously described^10^. Mice were placed in a closed Plexiglas chamber (40 × 40 × 30 cm) connected to a narrow corridor serving as a refuge zone. A ceiling-mounted LCD screen was used to present a looming visual stimulus consisting of a rapidly expanding dark circle simulating an approaching predator. Mice were allowed to freely explore the apparatus for 5 min before stimulus presentation. The looming stimulus was delivered in 2-3 trials per session, with inter-trial intervals of approximately 3 min. Behavior was recorded and analyzed offline by an experimenter blinded to group identity. The following parameters were quantified: (i) flight latency, defined as the time from stimulus onset to initiation of escape, and (ii) refuge duration, defined as the time spent in the refuge before re-emergence. Animals were randomly assigned to experimental groups, and all behavioral scoring was conducted under blinded conditions.

### Optogenetic manipulation

A blue laser (473 nm; Aurora-220-473, NEWDOON, Hangzhou, China) was used for optogenetic stimulation. Light was delivered through an implanted optical fiber at an intensity of 5 mW at the fiber tip, with stimulation frequencies of 1, 20, or 40 Hz (10 ms pulse width).

For closed-loop optogenetic activation during REM sleep, mice were first habituated to the EEG/EMG recording setup for 2 days. REM sleep onset was identified online based on low EMG tone and dominant theta-band (5–8 Hz) activity in the EEG. Light stimulation was manually triggered immediately upon REM onset and terminated upon transition to wakefulness or NREM sleep. For experiments assessing behavioral consequences, 40 Hz stimulation was applied selectively during REM sleep between 09:00 and 16:00. EEG/EMG recordings were discontinued after 16:00, followed immediately by open-field and looming behavioral testing.

To determine the state-dependent effects of GPe PV⁺ neuron activation, mice were recorded between 13:00 and 18:00. Experimenters continuously monitored EEG/EMG signals in real time and manually triggered stimulation upon detection of either NREM or REM sleep. During REM sleep, stimulation was initiated at REM onset and continued until REM termination. During NREM sleep, stimulation was initiated at NREM onset and maintained for 30 s. No-light, 1 Hz, 20 Hz, and 40 Hz stimulation conditions were delivered in a randomized order. Each mouse underwent 3–6 trials per condition, with an inter-trial interval of at least 3 min. Light delivery was synchronized with EEG/EMG recordings via transistor–transistor logic (TTL) signals. Sleep scoring and data analysis were performed by experimenters blinded to experimental conditions.

For optogenetic activation of microglia, blue light (470 nm) was delivered through the same implanted optical fiber in Cx3cr1-CreER; Ai27 mice, in which microglia express channelrhodopsin. Light stimulation (10 ms pulse width, 10 Hz, 5 mW at the fiber tip) was applied, and neuronal responses were simultaneously recorded using optrode configurations.

### Chemogenetic manipulation

For chemogenetic manipulation experiments, CPS was initiated two weeks after viral injection and EEG/EMG implantation. During CPS, mice were exposed to close physical contact with a rat for 10 min daily between 17:00 and 18:00. Clozapine-N-oxide (CNO; 1 mg/kg) was administered intraperitoneally at 08:00 and 13:00 on the following day. On CPS days 4, 8, and 12, EEG/EMG recordings were conducted during the stress paradigm to assess the effects of chemogenetic manipulation on REM sleep.

### Optrode recording

Signals were acquired using an Apollo II high-performance data acquisition system (Bio-Signal Technologies, China), amplified, high-pass filtered at 300 Hz with a third-order filter, and digitized at 30 kHz. Recordings were performed in freely behaving mice after at least two weeks of post-surgical recovery, including two days of habituation to the recording cables.

Spike sorting was conducted offline using a valley-seeking algorithm (Offline Sorter, Plexon). Single units were identified based on the presence of refractory periods, waveform quality, and principal component–based cluster isolation. Firing rate analyses were performed using NeuroExplorer 4 (Nex Technologies).

Optogenetic identification of GPe parvalbumin (PV) neurons was performed at the end of each recording session.^52^ Light pulse trains (635 nm, 10 s) at 20 or 40 Hz were delivered through the implanted optic fiber to activate ChrimsonR-expressing neurons under PV promoter control. Units were classified as optogenetically tagged if laser-evoked spikes met all of the following criteria: response reliability >0.7, first-spike latency <8 ms, jitter <3 ms, and high waveform similarity between laser-evoked and spontaneous spikes (correlation coefficient >0.9).

### Recording of splenic nerve activity

Recording of splenic nerve activity was performed under anesthesia, adapted from a previously described protocol.^53^ Mice were anesthetized with isoflurane, and the splenic nerve was surgically exposed using a standard approach. A custom cuff electrode was gently placed around the splenic nerve to allow stable nerve-electrode contact without stretching or compression. Neural signals were acquired using an Apollo II high-performance data acquisition system (Bio-Signal Technologies, China). Spontaneous splenic nerve activity was continuously recorded for 5 min under stable anesthetic conditions. For data analysis, recordings were band-pass filtered to reduce noise, and nerve firing activity was quantified offline. Average firing rates during the recording period were calculated for subsequent statistical analysis. For longitudinal assessment across CPS progression, splenic nerve activity was recorded in separate cohorts of mice at defined time points (CON, CPS day 1– 4), and firing rates were compared across groups.

### Splenic denervation

Splenic denervation was performed as previously described.^53^ Briefly, mice were anesthetized with isoflurane, and the spleen was exposed via a midline abdominal incision. Under a dissection microscope, absolute ethanol was repeatedly applied to the splenic vasculature using cotton applicators to ablate splenic nerve fibers running along the vessels, while carefully avoiding vascular damage. Sham-operated mice underwent identical surgical procedures except that saline was applied instead of ethanol. After surgery, mice were allowed to recover before subsequent experiments. The effectiveness of splenic denervation was verified by tyrosine hydroxylase (TH) immunofluorescence staining.

### Whole-cell slice electrophysiology

Whole-cell patch-clamp recordings were performed as previously described^54^ using glass micropipettes (3.7-4 MΩ) filled with an internal solution containing (in mM): 145 K-gluconate, 2 MgCl₂, 0.1 CaCl₂, 0.75 EGTA, 10 HEPES, 2 MgATP, and 0.3 Na₃GTP (pH 7.3, 290–295 mOsm). Signals were filtered at 1 kHz, digitized at 10 kHz, and acquired with an Axopatch 200B amplifier and Digidata 1440A system (Axon Instruments). Deep anesthesia was induced with 5% isoflurane prior to brain extraction. Coronal brain slices (250 μm) encompassing the GPe were prepared using a vibratome (Leica). Slices were initially allowed to recover for 30 min in ACSF maintained at 34 °C and subsequently kept at room temperature until recording. During electrophysiological recordings, slices were continuously superfused with ACSF at 30 °C at a flow rate of 2–3 mL/min. All recordings were completed within 4 hours after slice recovery. GPe parvalbumin (PV) neurons were identified by mCherry fluorescence following AAV9-PV-Cre and AAV9-hSyn-DIO-mCherry injection. For IL6 experiments, spontaneous firing activity was recorded under baseline conditions (10-15 min), followed by bath application of ACSF or IL-6 (10 ng/mL) for 10 min. Mean firing rates during the stable post-application period (5–10 min) were compared with baseline. In addition, neuronal excitability was assessed by injecting step currents (1000 ms duration) ranging from 0 to 200 pA to evoke action potential firing before and after IL6 incubation. Data were analyzed offline using Clampfit 10 (Molecular Devices).

### Immunofluorescent staining and imaging

For immunofluorescent staining of brain slices,^10^ mouse brains were fixed and obtained post cardiac perfusion with PBS and 4% PFA, and then dehydrated with gradient sucrose before being cut into 40 µm slices in the coronal plane around the injured site with a sliding microtome. The primary antibodies used in this study were: rabbit anti-Iba1 (Wako, 019-19741; Invitrogen, PA5-18039), rat anti-CD68 (BioLegend, 137002), rabbit anti-GFP (Rockland, 600-403-215), goat anti-GFP (Abcam, ab6673), rabbit anti-RFP (Rockland, 600-401-379), rabbit anti-IL6ST/gp130 (Bioss, bs-1459R), rabbit anti-CCR2 (Abcam, ab273050), rat anti-CD31 (BioLegend, 102501), rabbit anti-c-Fos (Cell Signaling Technology, #2250), mouse anti-tyrosine hydroxylase (TH; Merck Millipore, AB152), mouse anti-His-Tag (proteintech, 66005-1-Ig) and mouse anti-parvalbumin (PV; Millipore, MAB1572).

After washing, sections were incubated with species-appropriate fluorophore-conjugated secondary antibodies, including goat anti-mouse Alexa Fluor 488 (STARTER, S0B4017), goat anti-mouse Alexa Fluor 647 (Invitrogen, A21235), donkey anti-rat Alexa Fluor 488 (Abcam, ab150153), donkey anti-rat Alexa Fluor 594 (Abcam, ab105156), goat anti-rat Alexa Fluor 647 (STARTER, S0B4049), goat anti-rabbit Alexa Fluor 488 (STARTER, S0B4004), donkey anti-rabbit Alexa Fluor 594 (Jackson ImmunoResearch, 711-585-152), donkey anti-rabbit Alexa Fluor 647 (Abcam, ab150075), donkey anti-goat Alexa Fluor 488 (ABS, abs20026), and donkey anti-goat Alexa Fluor 555 (Invitrogen, A21432). Slices were sealed with fluorescent mounting medium containing DAPI (62248, Thermo Scientific), and imaged with a fluorescent scan microscope (FV3000, Olympus, Japan) or confocal laser microscope (LSM980, Zeiss, Germany).

Morphological and spatial analyses were performed using ImageJ. Iba1⁺ microglia were identified by threshold-based segmentation after background subtraction and counted using *Analyze Particles*. CD68 immunoreactivity was quantified as the CD68⁺ area within Iba1⁺ cells and normalized to the mean of the CON group. Microglial morphology was analyzed using binarization followed by *Skeletonize* and *Analyze Skeleton* to quantify branch number and process complexity. IL6ST expression was quantified as the IL6ST⁺ area relative to the EGFP⁺ area within the same field and normalized to the mean of the sgControl group. To assess monocyte–vascular distance, the minimum distance between CCR2⁺ cells and CD31⁺ blood vessels was calculated using the “Distance Map” and “Measure” functions after vessel segmentation.

### iDISCO lightsheet imaging

The 3D immunolabeling process of the whole mouse brain based on the iDISCO+ method was performed by Jarvis (Wuhan) biological pharmaceutical Co.,Ltd. Whole brain of CCR2^GFP^ mice were treated with gradient concentration methanol solutions for 3 days at room temperature and decolorized with 5% H2O2 solution overnight at 4℃. Then the sample was treated with PBST solution (0.2% TritonX-100 in PBS) for 3 days at room temperature with shaking. And the sample was immersed in a blocking solution (10% donkey serum in PTwH solution (0.2% Tween-20 in PBS)) for 3 days at room temperature and in the primary antibody solution (anti-GFP, dilution 1:500, A109002, Jarvis (Wuhan) biological pharmaceutical Co.,Ltd, China) for 6 days at room temperature. Then, the sample was washed for 2 days and was immersed in the second antibody solution (Alexa Fluor 488 Donkey anti-Rat IgG (H+L), 1:500 dilution, Thermofisher) for 6 days at room temperature. And the sample was washed for 2 days at room temperature. FDISCO (three-dimensional imaging of solvent-cleared organs with superior fluorescence preserving capability) tissue clearing reagent kit (JA11012, Jarvis (Wuhan) biological pharmaceutical Co.,Ltd, China) was used in this study. The clearing procedure was conducted entirely according to the kit instructions. To acquire images, the cleared samples were manually attached to the sample holder adapter. Subsequently, the samples were immersed in imaging reagent within a 3D printing sample chamber and excited with light sheets of different wavelengths. The resulting TIFF raw data images from light-sheet microscope were stitched and converted using LitScan software and subsequently reconstructed with Imaris software (Version 7.2.3, Bitplane, Switzerland).

### Tissue collection and blood sampling

Mice were deeply anesthetized with ketamine (100 mg/kg) and xylazine (10 mg/kg) and transcardially perfused with ice-cold phosphate-buffered saline (PBS). After perfusion, the skull was opened and the brain was removed. The choroid plexus was dissected under a stereomicroscope by gently isolating ventricular structures and collecting the choroid plexus tissue with fine forceps. For meningeal collection, the dura mater was carefully peeled from the inner surface of the skull, and the leptomeninges (arachnoid and pia mater) were separated from the cortical surface by gentle lifting and peeling with fine forceps to minimize parenchymal contamination. All tissues were immediately transferred to pre-chilled tubes for downstream analyses. Peripheral blood was collected into sodium heparin anticoagulant tubes to prevent clotting. Samples were kept on ice and processed promptly according to the requirements of downstream assays.

### Cells

Brain tissues were dissociated into single-cell suspensions using the Adult Brain Dissociation Kit (Miltenyi Biotec, 130-107-677) according to the manufacturer’s instructions, with gentle mechanical agitation using an incubator shaker during enzymatic digestion. After enzymatic and mechanical dissociation, myelin and cellular debris were removed using Debris Removal Solution, followed by erythrocyte lysis with Red Blood Cell Removal Solution (Miltenyi Biotec). PBMCs were isolated using a Mouse Peripheral Blood Lymphocyte Isolation Solution Kit (Solarbio Life Science, P8620) according to the manufacturer’s protocol. Spleens were minced in freshly prepared DMEM containing collagenase IV and DNase I and digested at 37 °C for 30 min with gentle agitation. Tissues were mechanically dissociated, filtered through a 70-μm strainer, and centrifuged. Red blood cells were lysed using red blood cell lysis buffer, followed by centrifugation, and the resulting single-cell suspensions were used for downstream analyses.

### Flow cytometry

CD45 antibody injection was performed as described.^18^ Briefly, five minutes before euthanasia, mice were intravenously injected with 2 μg of PE-conjugated anti-mouse CD45 antibody (clone 30-F11; BioLegend, #103105). Single-cell suspensions prepared from brain tissue and peripheral blood mononuclear cells (PBMCs) were subjected to flow cytometric analysis. Cells were first incubated with anti-CD16/32 antibody to block Fc receptors, followed by staining with a fixable viability dye (Zombie Aqua™, BioLegend, 423101). Cells were then stained with fluorophore-conjugated antibodies against surface markers, including anti-mouse CCR2 (BV605), Ly6C (BV650), CD45.2 (BV785), CD45 (PE), CD11b (PerCP), F4/80 (APC), Ly6G (APC-Cy7), CD3 (FITC), and CD19 (BV605). All antibodies were used at a 1:100 dilution, and staining was performed for 20 min at 4 °C. After staining, cells were washed with PBS, resuspended, and analyzed on a CytoFLEX SRT flow cytometer (Beckman Coulter). Flow cytometry data were analyzed using FlowJo software (Tree Star, USA). For downstream applications, including total RNA extraction, single-cell sequencing, and adoptive cell transfer, monocytes were enriched by fluorescence-activated cell sorting (FACS) based on CD45.2⁺CD11b⁺ expression using the CytoFLEX SRT (Beckman Coulter).

### Single cell RNA-seq and data analysis

Single cells isolated by fluorescence-activated cell sorting (BD FACS) were loaded onto Chromium microfluidic chips and processed using the Chromium Single Cell 3′ v3.1 chemistry on a 10x Chromium Controller (10x Genomics), following the manufacturer’s instructions. Cell concentration and viability were assessed using a TC20 automated cell counter (Bio-Rad), and approximately 8,000 viable cells (viability >90%) were used for library preparation. Single cells were encapsulated with Gel Beads-in-Emulsion (GEMs), followed by cell lysis and barcoded reverse transcription to generate full-length cDNA. After amplification, sequencing libraries were constructed using the Chromium Single Cell v3.1 Reagent Kit (10x Genomics). Libraries were sequenced on an Illumina NovaSeq 6000 platform according to standard protocols. Raw sequencing data were processed using the Cell Ranger pipeline (10x Genomics) with default parameters. Reads were demultiplexed, aligned to the mouse reference genome, and unique molecular identifiers (UMIs) were counted to generate gene– cell expression matrices. Downstream analyses were performed using Seurat. Cells with fewer than 200 detected genes and genes expressed in fewer than three cells were excluded, and low-quality or non-target cell populations were removed based on standard quality-control metrics. The expression matrices were analyzed using Seurat v3.1.5 in R. Data were log-normalized using the *NormalizeData* function with a scale factor of 10,000. The top 2,000 highly variable genes were identified using *FindVariableFeatures*, and the data were scaled using *ScaleData*. Principal component analysis was performed on the variable genes, and the number of principal components used for downstream analyses was determined using *ElbowPlot*. Cell clustering was performed using *FindNeighbors* and *FindClusters*, and two-dimensional visualization was generated using Uniform Manifold Approximation and Projection. Cluster identities were assigned based on the expression of canonical marker genes, with reference-based annotation using SingleR used as a secondary validation. Differential gene expression analysis between experimental conditions within each annotated cell population was performed using the *FindMarkers* function. Differentially expressed genes defined by adjusted P values were subjected to Gene Ontology enrichment analysis, which was performed separately for upregulated and downregulated gene sets. Enriched biological process terms were identified based on adjusted P values, and representative GO terms were selected for visualization.

### Human brain transcriptomic dataset analysis

Human GPe transcriptomic data were obtained and analyzed using the public web portal of the Human and Mammalian Brain Atlas. ^51^ Single-nucleus RNA-seq data from human GPe were analyzed to identify GABAergic neuronal populations, including SOX6⁺/CTXND1⁺ and LHX6⁺/LHX8⁺ subtypes. Gene expression levels of IL6ST, IL6R, and canonical neuronal markers were extracted using UMAP embeddings, which were visualized using the portal’s native plotting functions, without custom computational analysis.

### Bulk RNA sequencing

Tissues were rapidly dissected in ice-cold phosphate-buffered saline (PBS). Total RNA was extracted using TRIzol reagent (Invitrogen, CA, USA) according to the manufacturer’s instructions. RNA concentration and integrity were assessed using an Agilent 2100 Bioanalyzer (Agilent Technologies, Santa Clara, CA, USA), and samples meeting quality criteria were used for library preparation. RNA-seq libraries were constructed using the VAHTS Universal V10 RNA-seq Library Prep Kit (Premixed Version) following the manufacturer’s protocol. Library preparation, sequencing, and primary bioinformatic processing were performed by OE Biotech Co., Ltd. (Shanghai, China). For bulk RNA sequencing of splenic CD11b⁺ cells from Cx3cr1–Adrb2 conditional knockout mice, single-cell suspensions were prepared from spleens and CD11b⁺ cells were enriched by positive magnetic selection using the MACS system (Miltenyi Biotec, 130-126-725) prior to RNA extraction and sequencing.

### RNA purification and qPCR

Total RNA was isolated from brain tissue, peripheral blood mononuclear cells (PBMCs), and spleen using Freezol Reagent (Vazyme) according to the manufacturer’s instructions. Complementary DNA (cDNA) was synthesized using the Evo M-MLV RT Mix Kit with gDNA Clean for qPCR (Accurate Biology). Quantitative PCR (qPCR) was performed using 2× SYBR Green Premix (ROX Plus; Accurate Biology) in a final reaction volume of 20 μL on a QuantStudio 7 Flex Real-Time PCR System (Applied Biosystems). Amplification was carried out under the following conditions: 95 °C for 30 s, followed by 40 cycles of 95 °C for 5 s and 60 °C for 30 s. Relative mRNA expression levels were calculated using the comparative cycle threshold method (2^−ΔΔCt) and normalized to the housekeeping gene GAPDH.

Primers used in the study are as follows:

GAPDH: sense 5’ -CAGAACATCATCCCTGCATC-3’;

antisense 5’-TACTTGGCAGGTTTCTCCAG-3’;

IL6: sense 5’ACCGCTATGAAGTTCCTCTC-3’;

antisense 5’-CTCTGTGAAGTCTCCTCTCC-3’.

### Statistics

Data from all trials were averaged for each animal before cross-animal statistical analyses. All data are presented as mean ± SEM. Statistical analyses were performed using GraphPad Prism (GraphPad Software, Inc.). Normality was assessed using the Shapiro–Wilk test and homogeneity of variance was evaluated using the Brown–Forsythe test in GraphPad Prism. Depending on the experimental design, Mann-Whitney U test, Friedman test, Kruskal-Wallis test, one-way ANOVA, or two-way ANOVA were used as appropriate (Table S1). When ANOVA revealed significant main effects or interactions, post hoc multiple-comparison tests were performed as specified in Table S1. A P value < 0.05 was considered statistically significant. Statistical significance is indicated as follows: *P < 0.05, **P < 0.01, ***P < 0.001, ****P < 0.0001; ns, not significant.

---

**Extended Data Fig 1.**
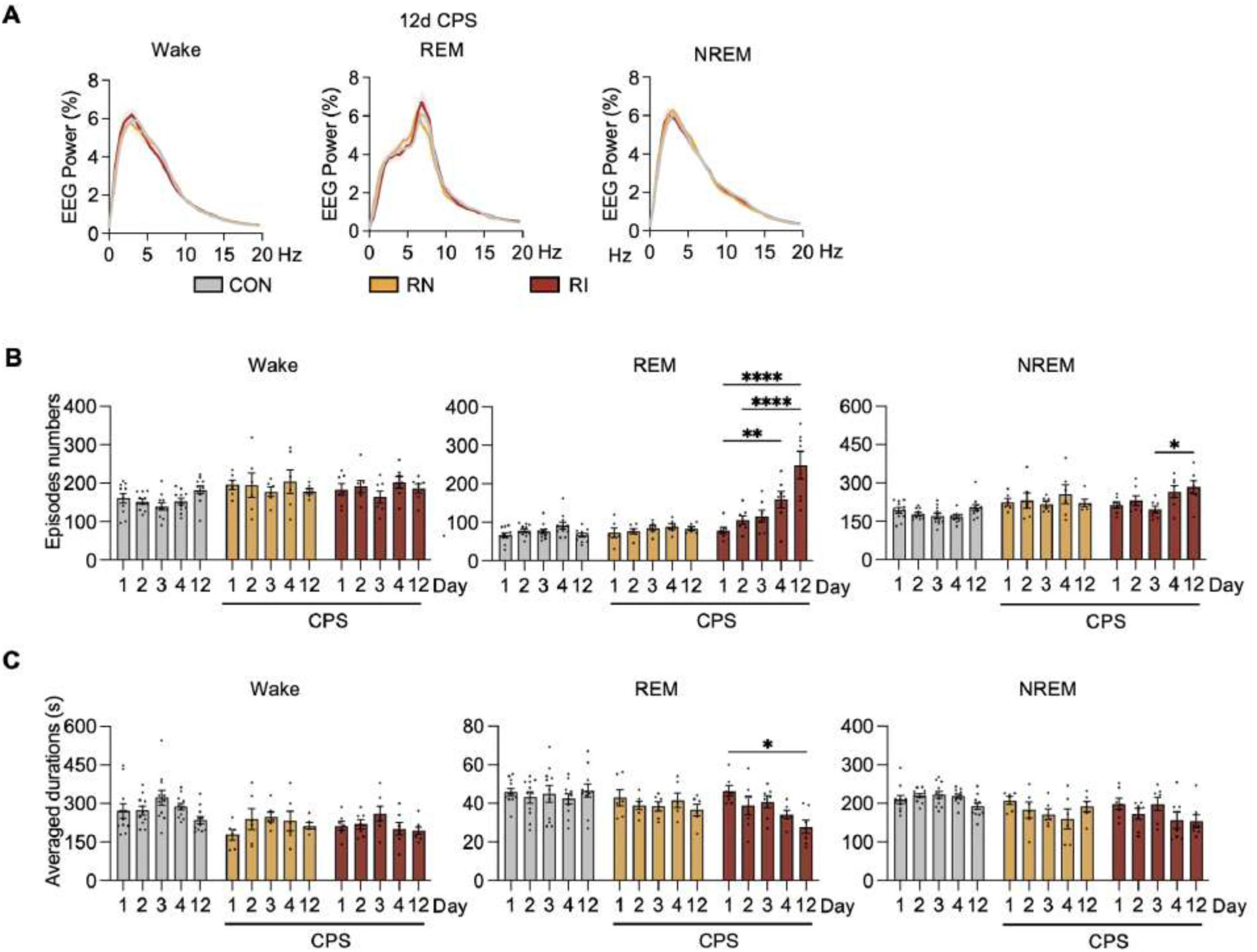
Temporal changes in sleep architecture under chronic predator stress (CPS), related to Figure 1. (A) EEG power spectra during each vigilance state during 24-hour recording on day 12 of CPS in CON, RN, and RI mice. (B) Number of episodes for wakefulness, REM sleep, and NREM during 24-hour recording across multiple CPS time points (Day 1, Day 2, Day 3, Day 4, and Day 12). (C) Average episode duration for wakefulness, REM sleep, and NREM sleep across CPS. All data are shown as mean ± SEM. Statistical significance is indicated as *P* < 0.05 (*), < 0.01 (**), < 0.001 (***), and < 0.0001 (****). See also Table S1.

**Extended Data Fig 2.**
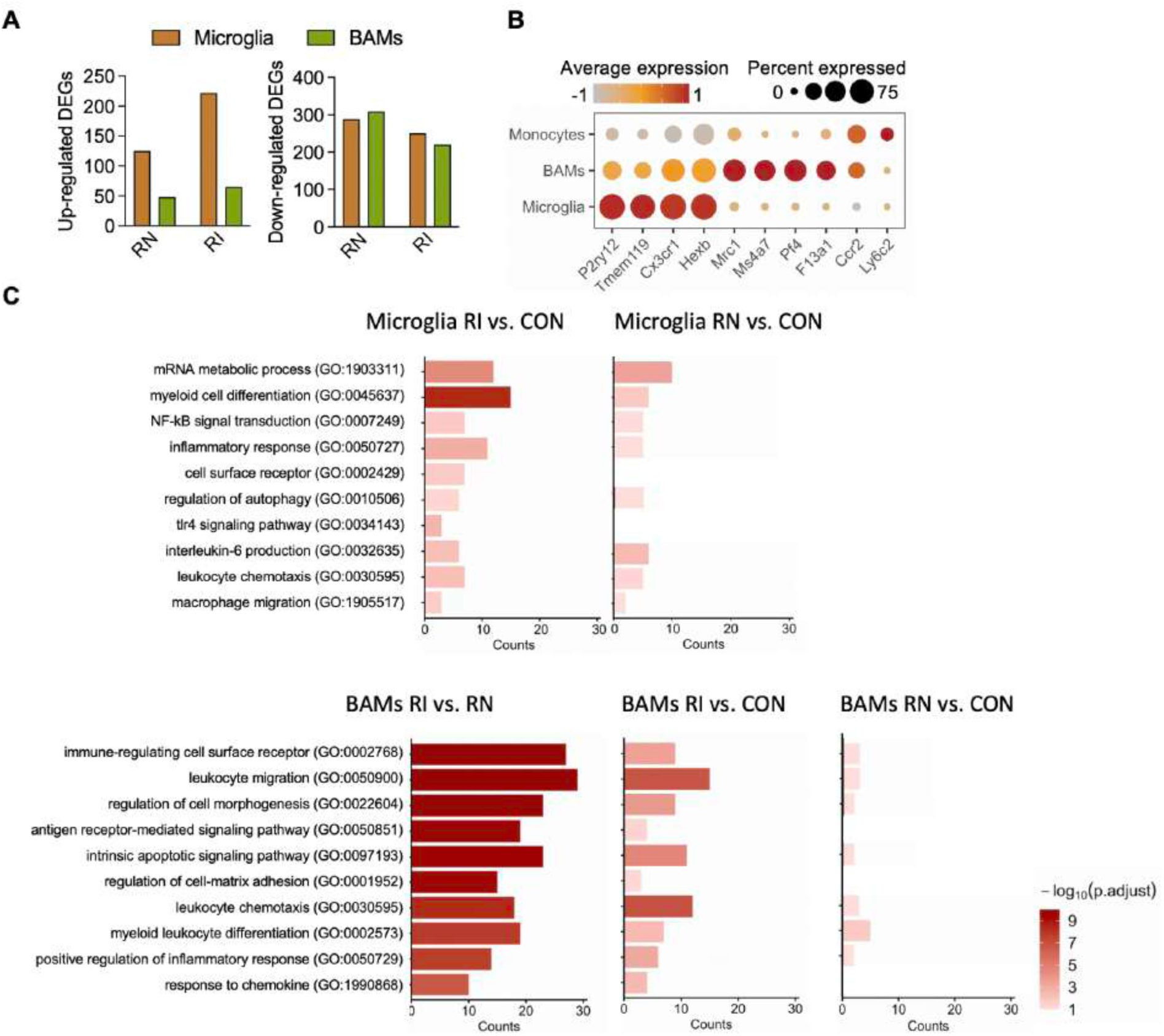
Differential transcriptional profiles of microglia and BAM populations under CPS, related to. Figure 1 (A) Numbers of upregulated and downregulated differentially expressed genes (DEGs) in microglia and BAMs from RN and RI mice compared with controls (CON). (B) Dot plot showing average expression levels and the proportion of cells expressing canonical marker genes across monocytes, BAMs, and microglia. Color intensity indicates scaled average expression, and dot size represents the percentage of expressing cells. (C) Gene Ontology (GO) enrichment analysis of biological pathways commonly upregulated in both RI versus RN and RI versus CON in microglia (top) and BAMs (bottom). Fewer pathways are observed in RN versus CON because some shared pathways are not enriched in this comparison. All data are shown as mean ± SEM. Statistical significance is indicated as *P* < 0.05 (*), < 0.01 (**), < 0.001 (***), and < 0.0001 (****). See also Table S1.

**Extended Data Fig 3.**
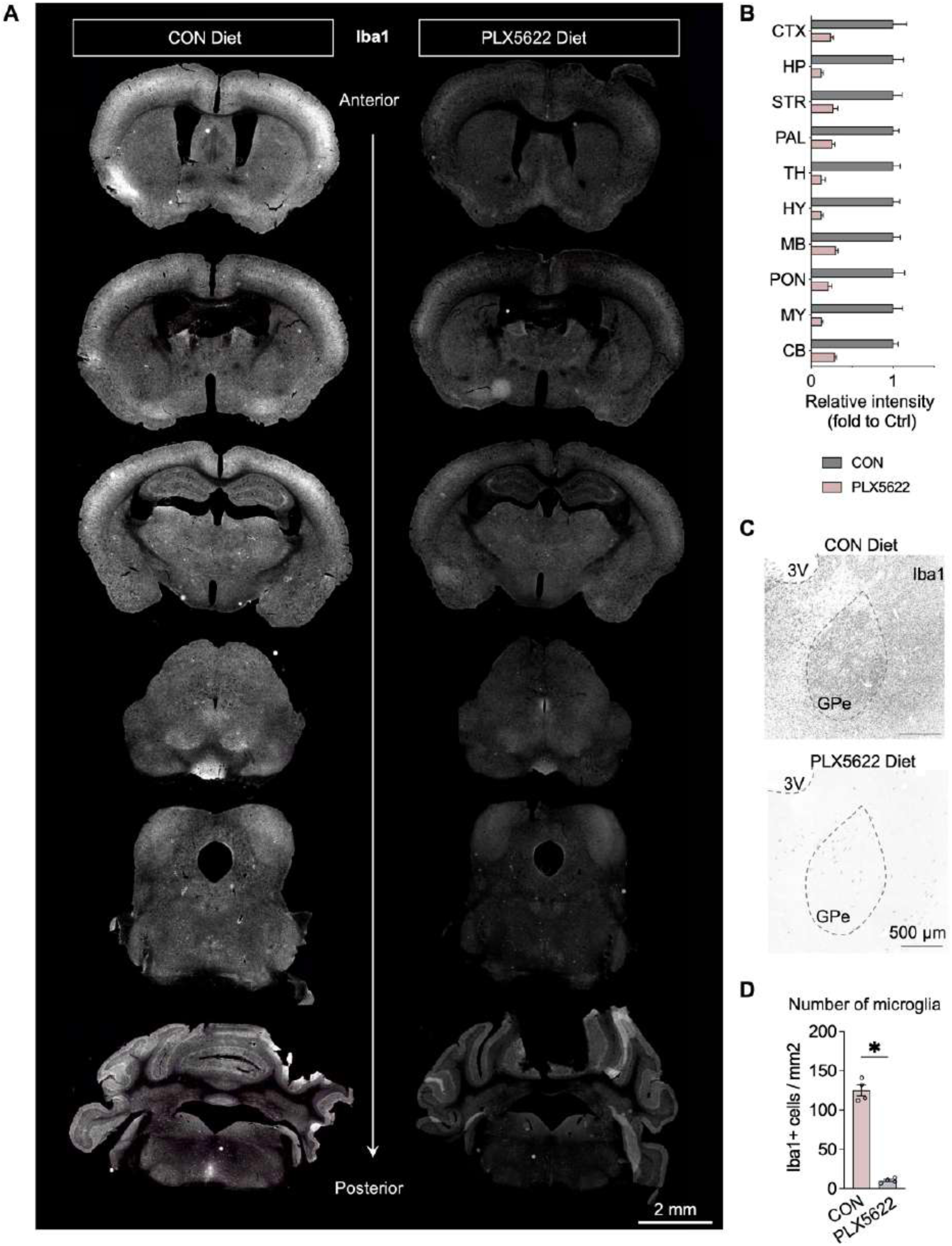
Effects of microglia depletion by PLX5622 under chronic predator stress, related to Figure 1. (A) Representative images showing Iba1⁺ cells across the brain in control (CON) and PLX5622-treated mice (scale bar, 2 mm). (B) Quantification of microglial density under CON and PLX5622 conditions. (C) Representative images of Iba1⁺ cells in the GPe from CON and PLX5622-treated mice (scale bar, 500 μm). (D) Quantification of microglial density in the GPe under CON and PLX5622 conditions. All data are shown as mean ± SEM. Statistical significance is indicated as *P* < 0.05 (*), < 0.01 (**), < 0.001 (***), and < 0.0001 (****). See also Table S1.

**Extended Data Fig 4.**
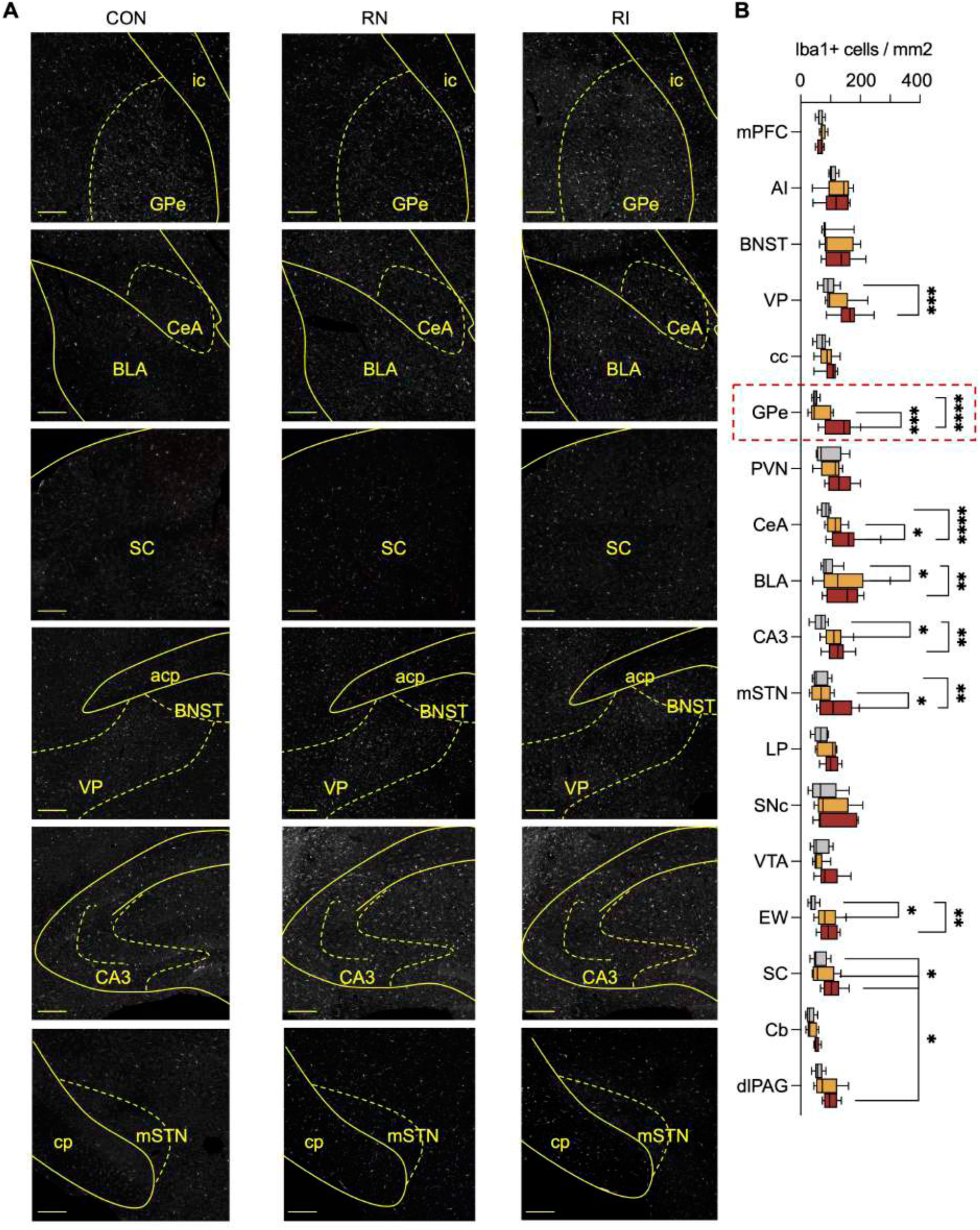
Iba1⁺ microglia across multiple brain regions following CPS, related to Figure 2. (A) Representative immunofluorescence images showing Iba1⁺ cells in multiple brain regions from CON, RN, and RI mice. Scale bar, 200 μm. (B) Quantification of Iba1⁺ cell density (cells per mm²) across multiple brain regions in CON, RN, and RI mice. Regions shown include the medial prefrontal cortex (mPFC), agranular insular cortex (AI), bed nucleus of the stria terminalis (BNST), ventral pallidum (VP), corpus callosum (cc), external globus pallidus (GPe), paraventricular nucleus of the hypothalamus (PVN), central nucleus of the amygdala (CeA), basolateral amygdala (BLA), Cornu Ammonis area 3 (CA3), medial subthalamic nucleus (mSTN), lateral posterior thalamic nucleus (LP), substantia nigra pars compacta (SNc), ventral tegmental area (VTA), Edinger–Westphal nucleus (EW), superior colliculus (SC), cerebellum (Cb), and dorsolateral periaqueductal gray (dlPAG). All data are shown as mean ± SEM. Statistical significance is indicated as P < 0.05 (*), < 0.01 (**), < 0.001 (***), and < 0.0001 (****). See Table S1 for details.

**Extended Data Fig 5.**
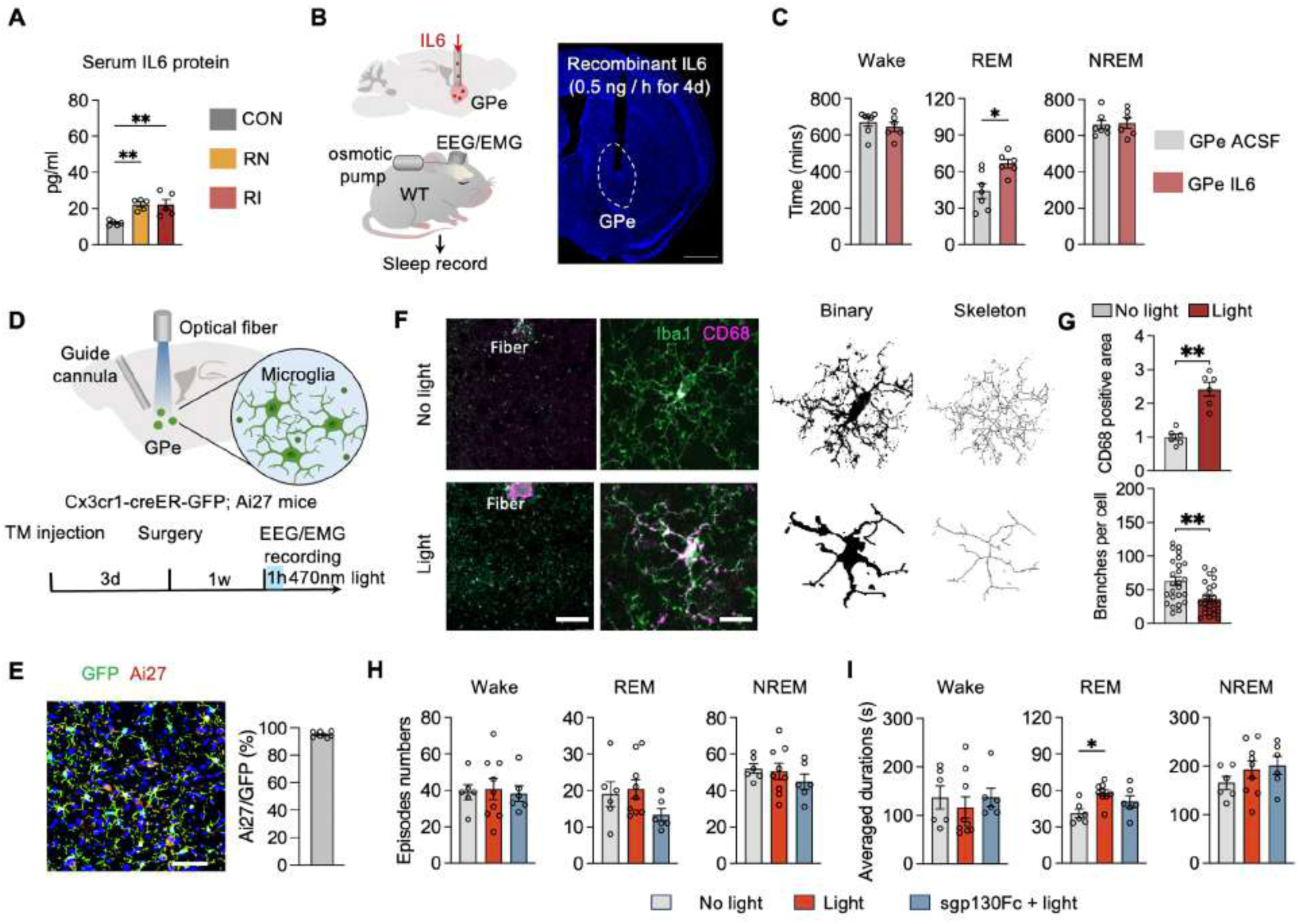
IL6 signaling in the GPe modulates REM sleep, related to Figure 2. (A) Serum IL6 protein levels measured across CON, RN, and RI groups. (B) Schematic and representative image of local IL6 delivery to the GPe using an osmotic pump. ACSF or recombinant IL6 (0.5 ng/h) was infused into the GPe for 4 days (scale bar, 1 mm). (C) Time spent in wakefulness, REM sleep, and NREM sleep during 24-h recording in ACSF- and IL6-treated mice. (D) Schematic of microglia-specific optogenetic activation and local sgp130Fc infusion in the GPe. In Cx3cr1-CreER-GFP; Ai27 mice, an optical fiber and guide cannula were implanted above the GPe. Vehicle or sgp130Fc was locally infused 30 min before 470-nm light stimulation. (E) Representative images and quantification showing co-expression of GFP and Ai27 (tdTomato) in microglia from Cx3cr1-CreER-GFP; Ai27 mice (scale bar, 50 μm). (F) Representative microglial morphology without (top) and with (bottom) optogenetic stimulation, with corresponding binary and skeletonized images. (G) Quantification of CD68-positive area and microglial branch number per cell (left scale bar = 100 μm; right scale bar = 10 μm). (H) Episode number and average duration of wakefulness, REM sleep, and NREM sleep under the indicated conditions. (I) Average duration of wakefulness, REM sleep, and NREM sleep in No light, Light, and sgp130Fc + Light conditions. All data are shown as mean ± SEM. Statistical significance is indicated as P < 0.05 (*), < 0.01 (**). See Table S1 for details.

**Extended Data Fig 6.**
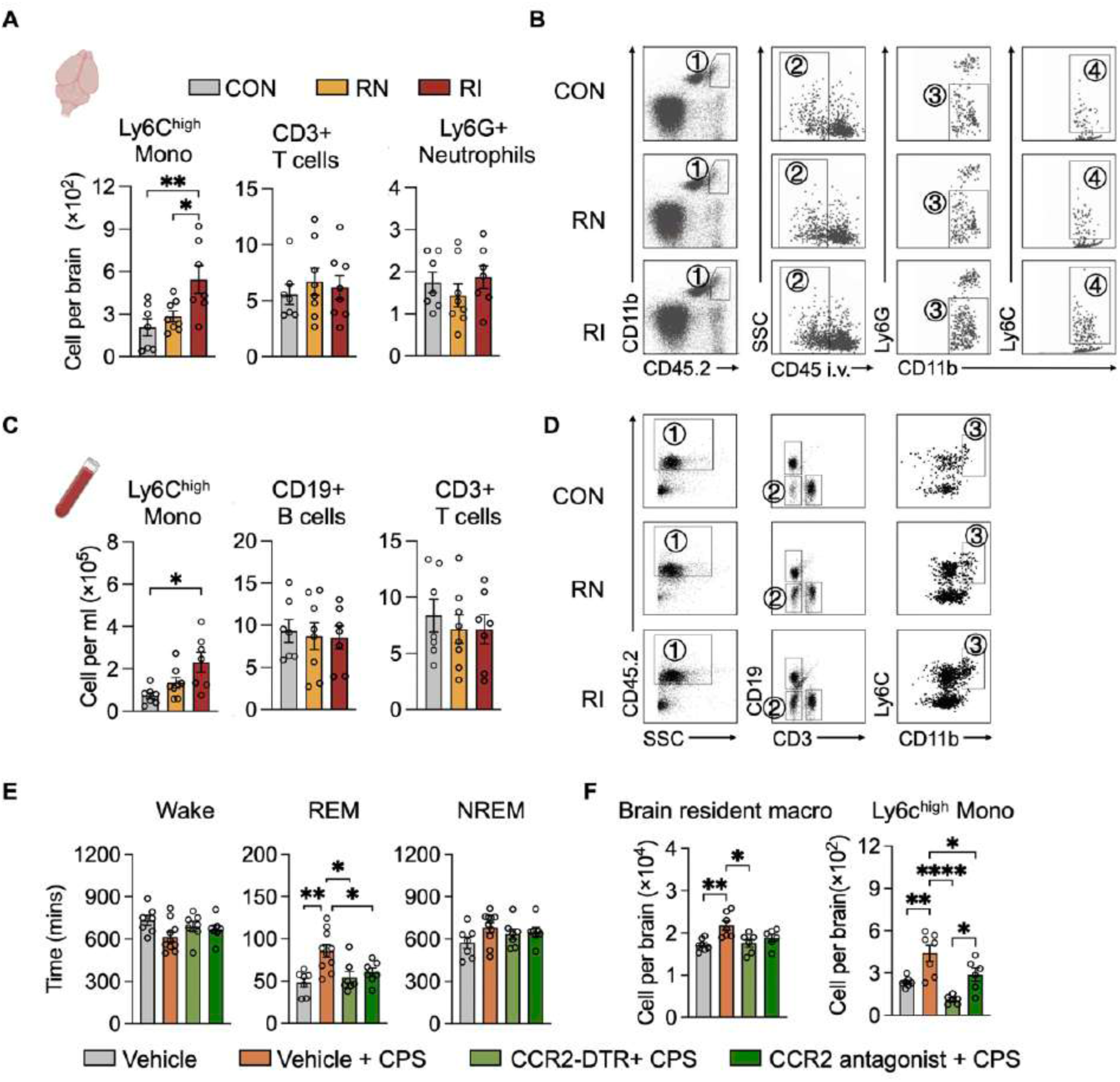
Altered immune cell composition in the brain and peripheral blood under CPS, related to Figure 3 (A) Quantification of Ly6C^high monocytes, CD3⁺ T cells, and Ly6G⁺ neutrophils in the brain across CON, RN, and RI mice. (B) Representative flow cytometry gating strategy for identification of infiltrating Ly6C^high monocytes based on CD11b, CD45, Ly6G, and Ly6C expression in CON, RN and RI mice. (C) Quantification of circulating Ly6C^high monocytes, CD19⁺ B cells, and CD3⁺ T cells in peripheral blood across experimental groups. (D) Representative flow cytometry plots illustrating peripheral blood immune cell populations in CON, RN, and RI mice. (E) Effects of CCR2-DTR–mediated monocyte depletion or pharmacological CCR2 antagonism on wakefulness, REM sleep, and NREM sleep following CPS. (F) Quantification of brain-resident macrophages and Ly6C^high monocytes per brain across experimental groups. All data are shown as mean ± SEM. Statistical significance is indicated as P < 0.05 (*), < 0.01 (**) and , < 0.0001 (****). See Table S1 for details.

**Extended Data Fig 7.**
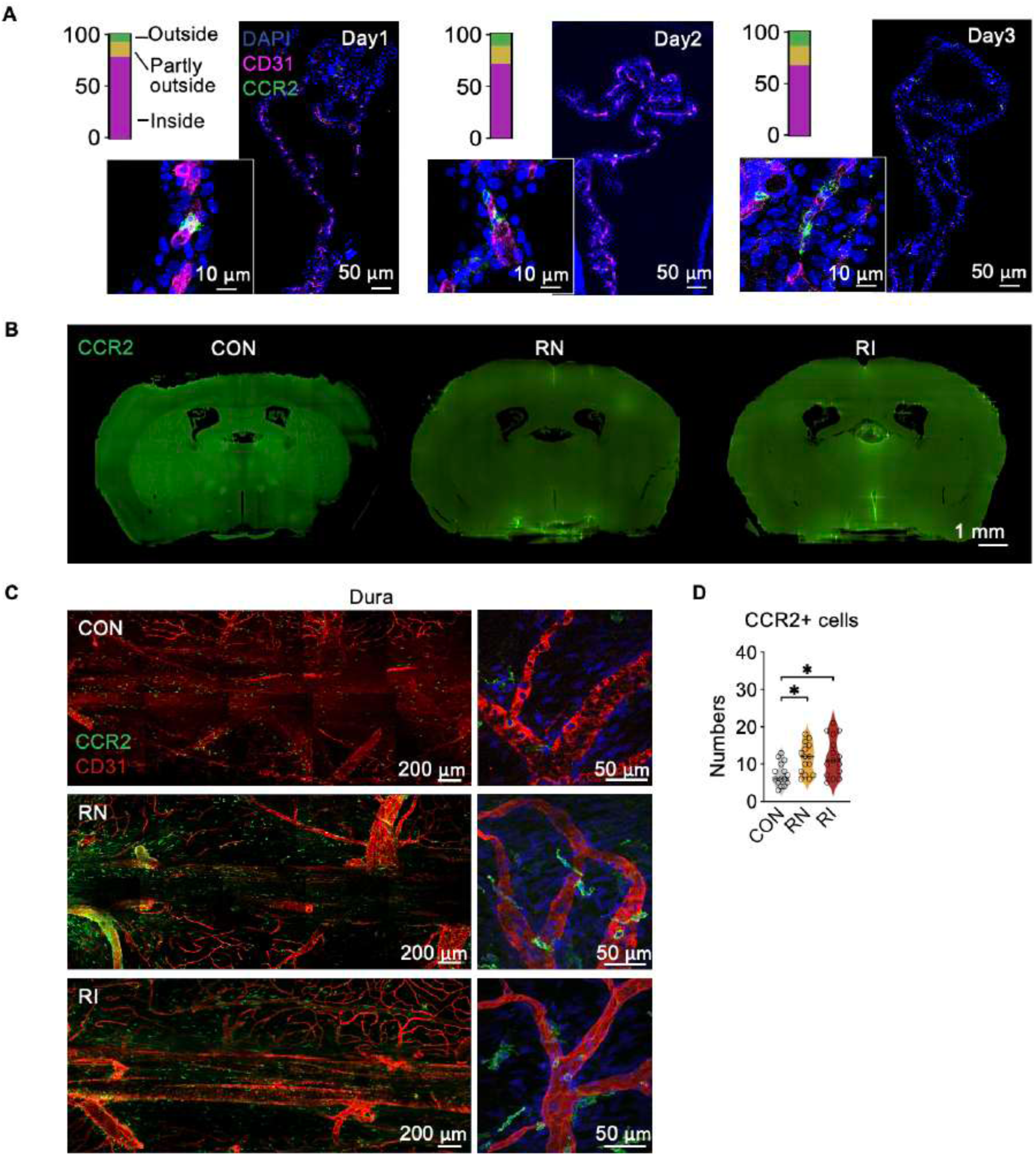
Quantification of CCR2⁺ cells in the choroid plexus (ChP) and meninges following CPS, related to Figure 3. (A) Immunofluorescence confocal images illustrating the spatial distribution of CCR2⁺ monocytes relative to ChP vasculature (CD31⁺) across successive days following CPS (Day1-Day3), insets show higher-magnification views. Stacked bar graphs indicate the proportion of CCR2⁺ cells located inside, partially outside, or outside the ChP boundary. (B) Representative coronal brain sections from whole-brain 3D imaging showing CCR2⁺ cell distribution in CCR2^GFP-DTR^ mice from CON, RN, and RI groups. (C) Representative immunofluorescence images of CCR2⁺ cells and CD31⁺ vasculature in the dura mater from CON, RN, and RI mice. (D) Quantification of CCR2⁺ cell numbers in the dura mater. All data are shown as mean ± SEM. Statistical significance is indicated as P < 0.05 (*). See Table S1 for details.

**Extended Data Fig 8.**
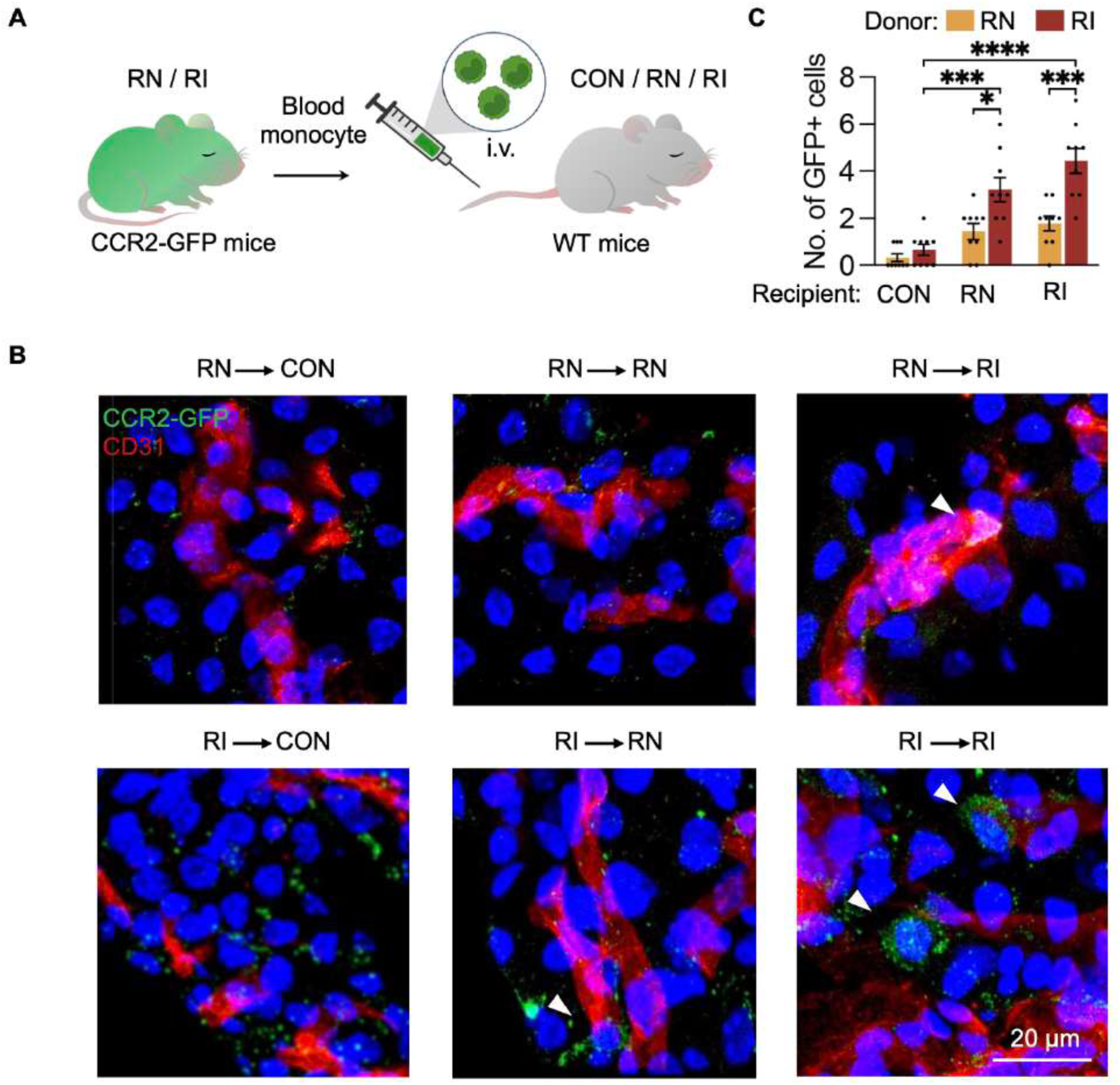
RI CCR2⁺ PBMC monocytes isolated from RI mice exhibited a greater capacity to infiltrate the ChP, related to Figure 3. (A) Peripheral blood CCR2⁺ monocytes were isolated from RN or RI CCR2^GFP-DTR^ donor mice and adoptively transferred into WT recipient mice via intravenous injection. Recipient mice were either exposed to CPS or left unexposed and were further classified into CON, RN, or RI groups based on their REM sleep phenotype. (B) Representative immunofluorescence images showing GFP⁺ CCR2⁺ donor-derived monocytes (green) localized near CD31⁺ vasculature (red) within the choroid plexus of recipient mice. Arrowheads indicate infiltrating GFP⁺ cells. (C) Quantification of GFP⁺ donor-derived cells detected in the choroid plexus of CON, RN, and RI recipient mice following transfer from RN or RI donors. All data are shown as mean ± SEM. Statistical significance is indicated as P < 0.05 (*), < 0.001 (***), and < 0.0001 (****). See Table S1 for details.

**Extended Data Fig 9.**
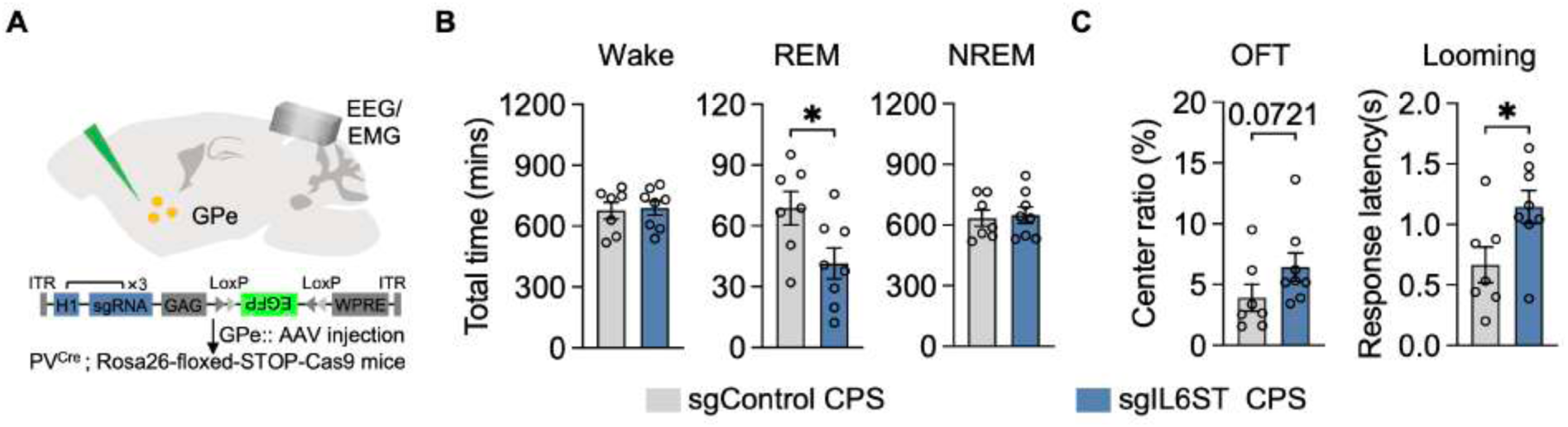
IL6ST signaling in GPe PV⁺ neurons is required for stress-induced REM sleep and threat-related behaviors, related to Figure 5. (A) Schematic of CRISPR-Cas9-mediated Il6st disruption in GPe PV⁺ neurons. sgControl denotes a non-targeting control sgRNA, whereas sgIL6ST targets Il6st for gene knockdown in PV⁺ neurons. (B) Total time spent in wakefulness, REM sleep, and NREM sleep over 24 h following CPS in sgControl and sgIL6ST mice. (C) Open-field test and looming assay performance in sgControl and sgIL6ST mice following CPS. All data are shown as mean ± SEM. Statistical significance is indicated as *p* < 0.05 (*), < 0.01 (**). See Table S1 for details.

**Extended Data Fig 10.**
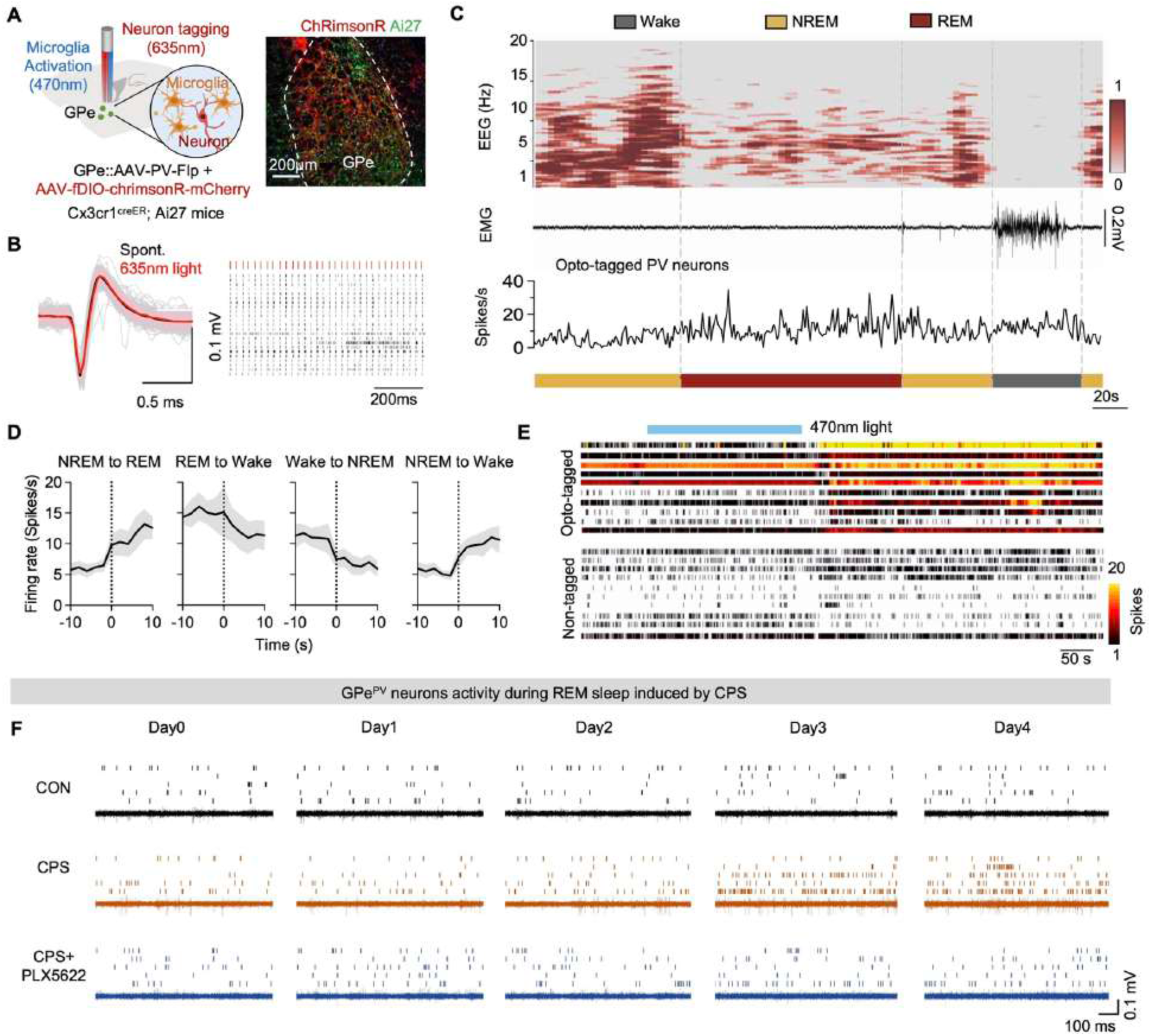
In vivo electrophysiological characterization of GPe PV neuron activity across CPS and microglial depletion, related to Figure 5. (A) Schematic of the experimental strategy combining optogenetic stimulation of Cx3cr1+ microglia and in vivo opto-electrophysiological recording of GPe PV+ neurons. Representative images show viral expression in the GPe of Cx3cr1-CreER; Ai27 mice. (B) Left, comparison of PV neuron firing evoked by 635-nm light stimulation versus spontaneous activity. Right, representative raster plot showing reliable spike responses to repeated 635-nm light pulses. (C) Representative simultaneous EEG, EMG, sleep-state scoring, and firing activity of GPe opto-tagged PV+ neurons. (D) Average firing rate dynamics of GPe opto-tagged PV+ neurons aligned to transitions between vigilance states. (E) Representative spike raster plots from opto-tagged and non-tagged neurons during microglial stimulation. (F) Representative spike raster plots and corresponding traces of GPe opto-tagged neurons recorded during REM sleep at baseline (Day 0) and at multiple time points following chronic predator stress (CPS; Day 1-4, and Day 12) in CON, CPS, and CPS + PLX5622-treated mice.

**Extended Data Fig 11.**
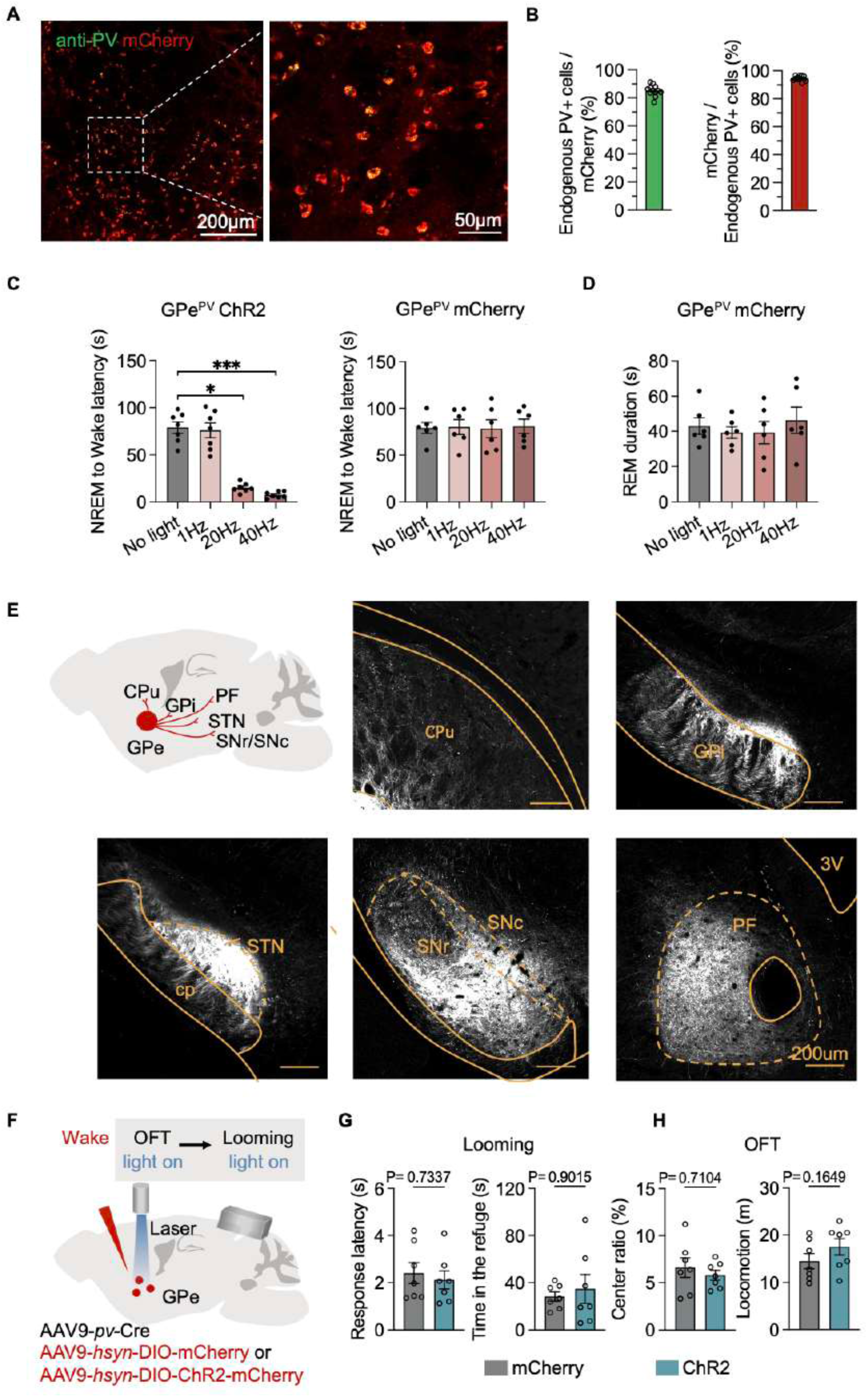
Validation and control experiments for optogenetic manipulation of GPe PV⁺ neurons, related to Figure 6. (A) Representative images showing mCherry expression and PV immunolabeling in the GPe following AAV9-PV-Cre–dependent viral expression, with a higher-magnification view of the boxed region (scale bar, 50 μm). (B) Quantification of the proportion of endogenous PV⁺ cells expressing mCherry and the proportion of mCherry⁺ cells that are PV⁺. (C) Effects of optogenetic stimulation of GPe^PV^ neurons expressing ChR2 on the latency of transitions from NREM sleep to wakefulness across different stimulation frequencies, with mCherry-expressing mice serving as controls. (D) REM sleep duration during optogenetic stimulation in GPe^PV^ mCherry control mice. (E) Schematic illustration of major downstream projection targets of GPe^PV^ neurons and representative images showing axonal projections in the caudate–putamen (CPu), subthalamic nucleus (STN), substantia nigra pars reticulata/compacta (SNr/SNc), globus pallidus internus (GPi), and parafascicular nucleus (PF) (scale bar, 200 μm). (F) Schematic of the optogenetic stimulation paradigm targeting GPe^PV^ neurons during looming and open-field test. (G) Effects of optogenetic activation of GPe^PV^ neurons during the looming task on looming-evoked defensive behavior. (H) Effects of optogenetic activation of GPe PV neurons during the open-field test. All data are shown as mean ± SEM. Statistical significance is indicated as P < 0.05 (*) and < 0.001 (***). See Table S1 for details.

## Notes

### Competing Interest Statement

The authors have declared no competing interest.

## References

1 Berger, M. & Riemann, D. Symposium: Normal and abnormal REM sleep regulation: REM sleep in depression-an overview. J Sleep Res 2, 211–223, doi:10.1111/j.1365-2869.1993.tb00092.x (1993).

2 Ross, R. J. et al. Rapid eye movement sleep disturbance in posttraumatic stress disorder. Biol Psychiatry 35, 195–202, doi:10.1016/0006-3223(94)91152-5 (1994).

3 Singareddy, R. K. & Balon, R. Sleep in posttraumatic stress disorder. Ann Clin Psychiatry 14, 183–190, doi:10.1023/a:1021190620773 (2002).

4 Freeman, D. et al. The effects of improving sleep on mental health (OASIS): a randomised controlled trial with mediation analysis. The Lancet Psychiatry 4, 749–758, doi:10.1016/S2215-0366(17)30328-0 (2017).

5 Freeman, D., Sheaves, B., Waite, F., Harvey, A. G. & Harrison, P. J. Sleep disturbance and psychiatric disorders. Lancet Psychiatry 7, 628–637, doi:10.1016/s2215-0366(20)30136-x (2020).

6 Krystal, A. D. Sleep therapeutics and neuropsychiatric illness. Neuropsychopharmacology 45, 166–175, doi:10.1038/s41386-019-0474-9 (2020).

7 Ford, D. E. & Kamerow, D. B. Epidemiologic study of sleep disturbances and psychiatric disorders. An opportunity for prevention? Jama 262, 1479–1484, doi:10.1001/jama.262.11.1479 (1989).

8 Howarth, N. E. & Miller, M. A. Sleep, Sleep Disorders, and Mental Health: A Narrative Review. Heart and Mind 8 (2024).

9 Harvey, A. G., Murray, G., Chandler, R. A. & Soehner, A. Sleep disturbance as transdiagnostic: consideration of neurobiological mechanisms. Clin Psychol Rev 31, 225–235, doi:10.1016/j.cpr.2010.04.003 (2011).

10 Tseng, Y.-T. et al. The subthalamic corticotropin-releasing hormone neurons mediate adaptive REM-sleep responses to threat. Neuron 110, 1223–1239.e1228, doi:10.1016/j.neuron.2021.12.033 (2022).

11 Schaefke, B., Li, J., Zhao, B., Wang, L. & Tseng, Y. T. Slumber under pressure: REM sleep and stress response. Prog Neurobiol 249, 102771, doi:10.1016/j.pneurobio.2025.102771 (2025).

12 Aime, M. et al. Paradoxical somatodendritic decoupling supports cortical plasticity during REM sleep. Science 376, 724–730, doi:10.1126/science.abk2734 (2022).

13 Wassing, R. et al. Restless REM Sleep Impedes Overnight Amygdala Adaptation. Curr Biol 29, 2351–2358.e2354, doi:10.1016/j.cub.2019.06.034 (2019).

14 Tseng, Y. T., Schaefke, B., Wei, P. & Wang, L. Defensive responses: behaviour, the brain and the body. Nat Rev Neurosci 24, 655–671, doi:10.1038/s41583-023-00736-3 (2023).

15 Tovote, P., Fadok, J. P. & Lüthi, A. Neuronal circuits for fear and anxiety. Nat Rev Neurosci 16, 317–331, doi:10.1038/nrn3945 (2015).

16 Fanselow, M. S. Negative valence systems: sustained threat and the predatory imminence continuum. Emerg Top Life Sci 6, 467–477, doi:10.1042/etls20220003 (2022).

17 Raison, C. L. & Miller, A. H. Malaise, melancholia and madness: the evolutionary legacy of an inflammatory bias. Brain Behav Immun 31, 1–8, doi:10.1016/j.bbi.2013.04.009 (2013).

18 Huynh, P. et al. Myocardial infarction augments sleep to limit cardiac inflammation and damage. Nature 635, 168–177, doi:10.1038/s41586-024-08100-w (2024).

19 Haykin, H. & Rolls, A. The neuroimmune response during stress: A physiological perspective. Immunity 54, 1933–1947, doi:10.1016/j.immuni.2021.08.023 (2021).

20 Frank, M. G., Baratta, M. V., Sprunger, D. B., Watkins, L. R. & Maier, S. F. Microglia serve as a neuroimmune substrate for stress-induced potentiation of CNS pro-inflammatory cytokine responses. Brain Behav Immun 21, 47–59, doi:10.1016/j.bbi.2006.03.005 (2007).

21 Xia, M. et al. Elevated IL-22 as a result of stress-induced gut leakage suppresses septal neuron activation to ameliorate anxiety-like behavior. Immunity 58, 218–231.e212, doi:10.1016/j.immuni.2024.11.008 (2025).

22 Cathomas, F. et al. Circulating myeloid-derived MMP8 in stress susceptibility and depression. Nature 626, 1108–1115, doi:10.1038/s41586-023-07015-2 (2024).

23 Wohleb, E. S., Powell, N. D., Godbout, J. P. & Sheridan, J. F. Stress-induced recruitment of bone marrow-derived monocytes to the brain promotes anxiety-like behavior. J Neurosci 33, 13820–13833, doi:10.1523/jneurosci.1671-13.2013 (2013).

24 Chan, K. L., Poller, W. C., Swirski, F. K. & Russo, S. J. Central regulation of stress-evoked peripheral immune responses. Nat Rev Neurosci 24, 591–604, doi:10.1038/s41583-023-00729-2 (2023).

25 Wohleb, E. S., McKim, D. B., Sheridan, J. F. & Godbout, J. P. Monocyte trafficking to the brain with stress and inflammation: a novel axis of immune-to-brain communication that influences mood and behavior. Front Neurosci 8, 447, doi:10.3389/fnins.2014.00447 (2014).

26 Lee, J. & Wheeler, M. A. Immune-brain plasticity underpins stress and affective behaviors. Neuron 114, 9–32, doi:10.1016/j.neuron.2025.11.020 (2026).

27 Jahromi, G. G. & Rezaei, N. Connecting the Dots: NLRP3 Inflammasome as a Key Mediator in the Intersection of Depression and Cardiovascular Disease – A Narrative Review. Heart and Mind 9 (2025).

28 Chung, E. N. et al. Psychedelic control of neuroimmune interactions governing fear. Nature 641, 1276–1286, doi:10.1038/s41586-025-08880-9 (2025).

29 Huang, S. et al. Disruption of the Na(+)/K(+)-ATPase-purinergic P2X7 receptor complex in microglia promotes stress-induced anxiety. Immunity 57, 495–512.e411, doi:10.1016/j.immuni.2024.01.018 (2024).

30 Ma, C. et al. Microglia regulate sleep through calcium-dependent modulation of norepinephrine transmission. Nature Neuroscience 27, 249–258, doi:10.1038/s41593-023-01548-5 (2024).

31 Tillmon, H. et al. Complement and microglia activation mediate stress-induced synapse loss in layer 2/3 of the medial prefrontal cortex in male mice. Nat Commun 15, 9803, doi:10.1038/s41467-024-54007-5 (2024).

32 Wu, J. et al. Integrating spatial and single-nucleus transcriptomic data elucidates microglial-specific responses in female cynomolgus macaques with depressive-like behaviors. Nature Neuroscience 26, 1352–1364, doi:10.1038/s41593-023-01379-4 (2023).

33 Reader, B. F. et al. Peripheral and central effects of repeated social defeat stress: monocyte trafficking, microglial activation, and anxiety. Neuroscience 289, 429–442, doi:10.1016/j.neuroscience.2015.01.001 (2015).

34 Wohleb, E. S. et al. Re-establishment of anxiety in stress-sensitized mice is caused by monocyte trafficking from the spleen to the brain. Biol Psychiatry 75, 970–981, doi:10.1016/j.biopsych.2013.11.029 (2014).

35 McKim, D. B. et al. Sympathetic Release of Splenic Monocytes Promotes Recurring Anxiety Following Repeated Social Defeat. Biol Psychiatry 79, 803–813, doi:10.1016/j.biopsych.2015.07.010 (2016).

36 Tseng, Y. T. et al. Systematic evaluation of a predator stress model of depression in mice using a hierarchical 3D-motion learning framework. Transl Psychiatry 13, 178, doi:10.1038/s41398-023-02481-8 (2023).

37 Yu, X. et al. A specific circuit in the midbrain detects stress and induces restorative sleep. Science 377, 63–72, doi:10.1126/science.abn0853 (2022).

38 Zhao, B. et al. An escape-enhancing circuit involving subthalamic CRH neurons mediates stress-induced anhedonia in mice. Neurobiol Dis 200, 106649, doi:10.1016/j.nbd.2024.106649 (2024).

39 Tynan, R. J. et al. Chronic stress alters the density and morphology of microglia in a subset of stress-responsive brain regions. Brain Behav Immun 24, 1058–1068, doi:10.1016/j.bbi.2010.02.001 (2010).

40 Cao, K. et al. Microglia modulate general anesthesia through P2Y(12) receptor. Curr Biol 33, 2187–2200.e2186, doi:10.1016/j.cub.2023.04.047 (2023).

41 Stranahan, A. M., Tabet, A. & Anikeeva, P. Region-specific targeting of microglia in vivo using direct delivery of tamoxifen metabolites via microfluidic polymer fibers. Brain Behav Immun 115, 131–142, doi:10.1016/j.bbi.2023.09.021 (2024).

42 Lv, Z. et al. Clearance of β-amyloid and synapses by the optogenetic depolarization of microglia is complement selective. Neuron 112, 740–754.e747, 10.1016/j.neuron.2023.12.003 (2024).

43 Kim, K. et al. Meningeal lymphatics-microglia axis regulates synaptic physiology. Cell 188, 2705–2719.e2723, doi:10.1016/j.cell.2025.02.022 (2025).

44 Qing, H. et al. Origin and Function of Stress-Induced IL-6 in Murine Models. Cell 182, 372–387.e314, doi:10.1016/j.cell.2020.05.054 (2020).

45 Osimo, E. F. et al. Inflammatory markers in depression: A meta-analysis of mean differences and variability in 5,166 patients and 5,083 controls. Brain Behav Immun 87, 901–909, doi:10.1016/j.bbi.2020.02.010 (2020).

46 Rose-John, S. Interleukin-6 signalling in health and disease. F1000R*es* **9**, doi:10.12688/f1000research.26058.1 (2020).

47 Zhu, X. A. et al. A neuroimmune circuit mediates cancer cachexia-associated apathy. Science 388, eadm8857, doi:10.1126/science.adm8857 (2025).

48 Leunig, A., Gianeselli, M., Russo, S. J. & Swirski, F. K. Connection and communication between the nervous and immune systems. Nat Rev Immunol 25, 912–933, doi:10.1038/s41577-025-01199-6 (2025).

49 Biltz, R. G., Sawicki, C. M., Sheridan, J. F. & Godbout, J. P. The neuroimmunology of social-stress-induced sensitization. Nat Immunol 23, 1527–1535, doi:10.1038/s41590-022-01321-z (2022).

50 Courtney, C. D., Pamukcu, A. & Chan, C. S. Cell and circuit complexity of the external globus pallidus. Nat Neurosci 26, 1147–1159, doi:10.1038/s41593-023-01368-7 (2023).

51 Hewitt, M., et al. A cross-species spatial transcriptomic atlas of the human and non-human primate basal ganglia. (2025).

52 Ding, X. et al. Neuroendocrine circuit for sleep-dependent growth hormone release. Cell 188, 4968–4979.e4912, doi:10.1016/j.cell.2025.05.039 (2025).

53 Zhang, X. et al. Brain control of humoral immune responses amenable to behavioural modulation. Nature 581, 204–208, doi:10.1038/s41586-020-2235-7 (2020).

54 Liu, X. A. et al. Interleukin 13 signaling modulates dopaminergic functions and nicotine reward in rodents. Mol Psychiatry 31, 622–634, doi:10.1038/s41380-025-03137-3 (2026).

